# Cryo-electron tomography reveals near-native archaeal chromatin architecture

**DOI:** 10.64898/2026.06.18.722658

**Authors:** Maximilian Dreimann, Fredrika Rajer, Quentin Durieux-Trouilleton, Tiana S. Behr, Sebastian Unger, Shingo Yamamoto, Naomichi Takemata, Haruyuki Atomi, Svetlana O. Dodonova

## Abstract

Most archaea package their genomes using histones, the evolutionary ancestors of eukaryotic chromatin proteins. Yet how archaeal chromatin is organized in cells has remained unknown. Here we provide the first high-resolution view of archaeal chromatin in a near-native state. We combine cryo-electron tomography, subtomogram averaging and single-particle cryo-electron microscopy to study chromatin released from gently lysed cells of the model archaeon *Thermococcus kodakarensis*. We show that archaeal chromatin is built from closed hypernucleosomes of variable length and determine structures of multiple hypernucleosome species, revealing a variable-beads-on-a-string architecture with flexible local contact geometries. We further show that chromatin organization changes with cellular state: logarithmic-phase cells are enriched in short hypernucleosomes, whereas stationary-phase cells form longer assemblies. Histone HTkB deletion shifts chromatin towards shorter hypernucleosomes in both growth phases and reduces local contact order. These results establish archaeal chromatin as a dynamic and complex system, shaped by growth phase and histone composition.

## Introduction

Archaea and eukaryotes use histone proteins to organize their genomic DNA. In archaea, histones form homodimers and heterodimers that can further oligomerize into extended assemblies, wrapping DNA into continuous superhelical structures termed hypernucleosomes, with each dimer binding ~30 bp of DNA^1^. This arrangement has been proposed to give rise to a variable “beads-on-a-string” architecture^2^, in which nucleosome size is not fixed but instead increases incrementally with the number of stacked dimers. In contrast, the fundamental unit of eukaryotic chromatin is the well-defined nucleosome core particle, consisting of a histone octamer composed of H2A, H2B, H3, and H4 core histones, that together wrap ~147 bp of DNA^3^.

Evidence for this model of archaeal chromatin organization first came from micrococcal nuclease (MNase) digestion experiments^2,4,5^, which revealed ladder-like patterns with a periodicity of ~30 bp rather than the single ~150 bp protected fragment characteristic of eukaryotic nucleosomes. Most studies of archaeal chromatin have focused on histones from the model euryarchaea *Thermococcus kodakarensis* and *Methanothermus fervidus*^*6–9*^. In these and many other hyperthermophilic archaea, histones can constitute a substantial fraction of the total proteome^10^, contributing not only to DNA stabilization under extreme conditions but also, potentially, to gene regulation^11^.

Structural and biophysical insights into archaeal chromatin have so far come almost exclusively from in vitro reconstitution studies based on histones from *T. kodakarensis* and *M. fervidus*^*6–8,12–14*^. Notably, many archaeal species encode two or more histone paralogs, for example, HTkA and HTkB in *T. kodakarensis* or HMfA and HMfB in *M. fervidus*. In addition to these canonical “nucleosomal” histones, some archaea also encode non-canonical histone variants, such as HTkC, where function remains less well understood^15^. Previous studies have shown that simultaneous deletion of both canonical histones is not possible in *T. kodakarensis*, whereas strains lacking either *htkA* or *htkB* genes individually remain viable^11^. A crystallographic study demonstrated that the HMfB histone from *M. fervidus* forms a minimal ~90 bp nucleosome that assembles into an extended hypernucleosome through crystal packing interactions^1^. Subsequent work on the HTkA histone from *T. kodakarensis* revealed two conformations in vitro: a superhelical compact “closed” architecture similar to HMfB, and a “slinky”-like arrangement in which two of such closed nucleosome units are oriented at ~90° angle to one another^16^. More recently, in vitro cryo-electron microscopy (cryo-EM) analysis showed that the Asgard archaeal HHoB histone forms extended hypernucleosomes that adopt both open and closed conformations (no slinkie arrangements were observed), expanding the range of architectures known for archaeal chromatin^17^.

Despite these advances, a major gap remains in our understanding of archaeal chromatin, as all structural information currently derives from in vitro systems, which may be strongly influenced by DNA sequence and length, protein purification strategies, and buffer conditions. In contrast, insights from native cellular material are limited to low-resolution bulk approaches such as micrococcal nuclease (MNase) digestion^2,18^ and a small number of AFM studies^19,20^. As a result, several fundamental questions remain unresolved: what is the native architecture of archaeal chromatin, which assemblies are most abundant in cells, do “slinky” arrangements exist in vivo, and how do histone variants and growth state influence chromatin organization?

Here, we provide the first high-resolution structural analysis of archaeal chromatin isolated directly from cells. By combining cryo-electron tomography (cryo-ET), subtomogram averaging (STA), and single-particle (SPA) cryo-EM, we analyze chromatin in crude lysates of *T. kodakarensis* without purification or in vitro reconstitution. At high resolution, we resolve multiple hypernucleosome species of different lengths, quantify their distribution, and define the geometric relationships between neighboring particles within chromatin fibers. Our results reveal that archaeal chromatin adopts a variable-beads-on-a-string architecture in which closed hypernucleosomes form the fundamental structural units. We further show that chromatin organization depends on both growth phase and histone composition: logarithmic-phase cells are enriched in shorter hypernucleosomes, whereas stationary-phase cells accumulate longer species, and deletion of *htkB* gene strongly shifts chromatin toward shorter assemblies. Together, these findings establish a structural framework for understanding archaeal chromatin organization in near-native material and reveal how chromatin architecture is shaped by physiological state and histone composition.

## Results

### Cryo-ET reveals a variable-beads-on-a-string organization of near-native archaeal chromatin

To investigate chromatin organization in *T. kodakarensis*, we used our laboratory strain KU216, a uracil-auxotrophic strain generated from the wild-type *T. kodakarensis* strain KOD1^21^. Cells were grown in nutrient-rich medium containing 0.8 × concentration of Artificial Sea Water (ASW) for 24 h at 85 °C under anaerobic conditions to stationary phase (Fig. 1a). Cells were then gently lysed by resuspension in 0.4 × ASW supplemented with 0.05% (v/v) Triton X-100 and 10 µg/mL RNase A, immediately applied to EM grids, and plunge-frozen (Methods, Fig. 1a). During data collection, regions adjacent to visible cell debris were targeted for tilt-series acquisition.

**Fig. 1.**
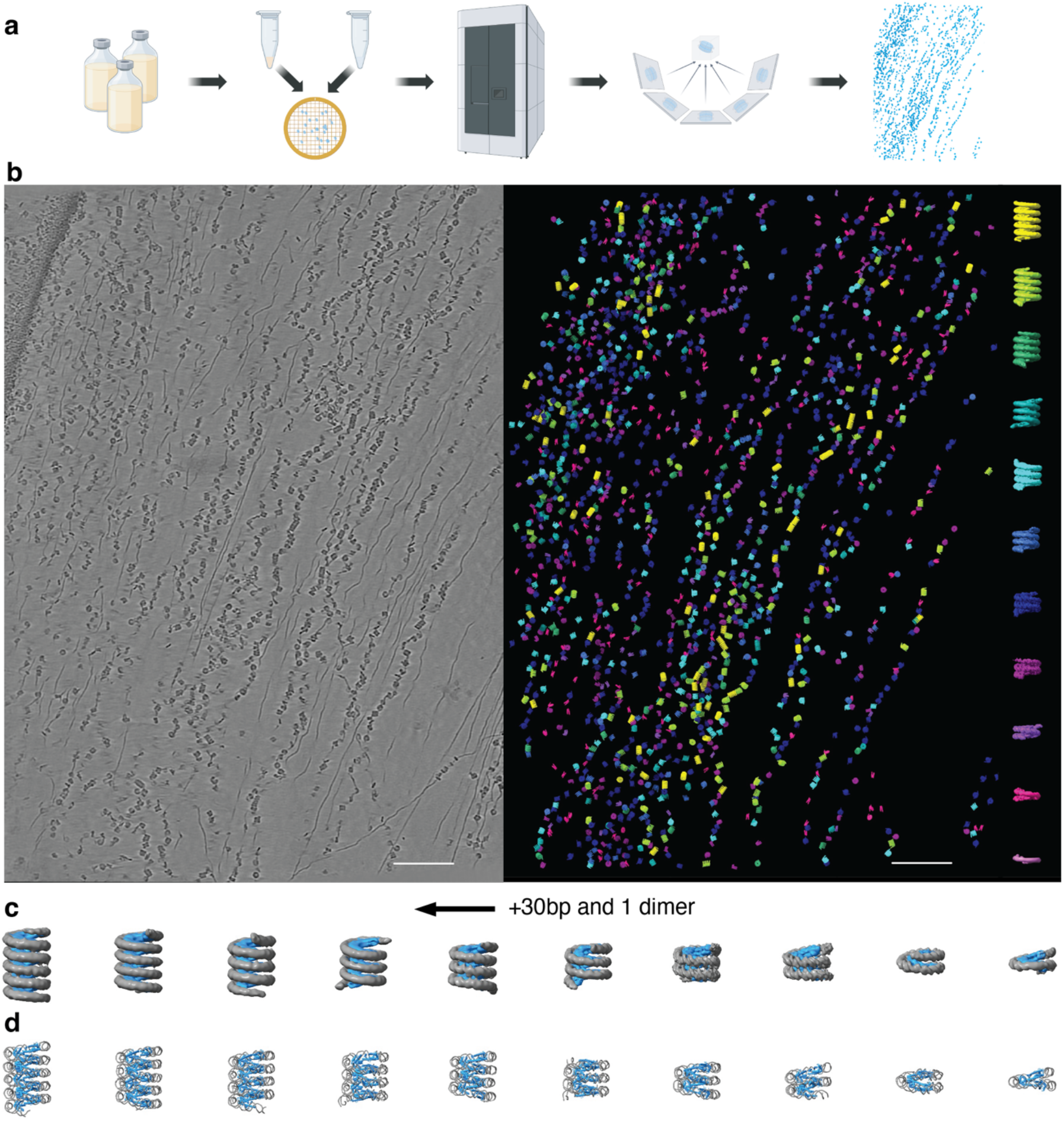
Cryo-ET of *T. kodakarensis* chromatin from stationary-phase cells. a. Schematic workflow of cryo-ET sample preparation, data acquisition, and analysis. b. Representative slice through a tomogram of *T. kodakarensis* chromatin. Left: raw tomographic slice; right: subtomograms pasted back into the tomogram at their refined positions and orientations. Scale bar - 100 nm. c. Ten STA maps at resolutions of 6.5-11.7 Å, with DNA shown in grey and histones in blue. d. Corresponding structural models of the different hypernucleosomal classes, with DNA shown in grey and histones in blue.

Visual inspection of reconstructed tomograms revealed long chromatin fibers composed of multiple hypernucleosomes of different sizes interspersed with free DNA regions (Fig. 1b). In many fibers, hypernucleosomes were positioned nearly continuously, with minimal linker DNA between neighboring particles, whereas other regions contained longer stretches of free DNA.

Tomograms containing chromatin fibers were then used for particle picking and STA (see Methods, Supp. Fig. 1), yielding 10 classes of hypernucleosome particles of varying length at resolutions of up to 6.5 Å (range, 6.5–11.7 Å) (Fig. 1c; Supp. Fig. 1). Corresponding models were generated for each hypernucleosomal species (Fig.1d, see Methods). This analysis showed that assemblies containing between 3 and ~12 histone dimers, corresponding to ~90 - 360 bp of wrapped DNA, represent the most abundant hypernucleosome species in stationary-phase *T. kodakarensis* chromatin ex situ (Fig. 1 c, d).

Mapping the subtomogram averages back into the tomograms on the basis of their positions and orientations revealed a variable-beads-on-a-string organization of long archaeal chromatin fibers (Fig. 1b-d, Video S1). Hypernucleosomes of different sizes formed irregular fibers separated by variable lengths of free DNA, generating local regions of densely packed chromatin interspersed with more accessible segments. The size distribution of hypernucleosomes observed in these tomograms is consistent with earlier MNase digestion experiments, which likewise showed a broad distribution of protected DNA fragments (~60-500 bp)^2^.

### Deletion of histone HTkB gene leads to loss of extended hypernucleosomes

To examine the contribution of histone HTkB to chromatin organization, we generated a *T. kodakarensis* KU216 derivative strain lacking *htkB* gene (hereafter termed *ΔhtkB*) essentially as described in ^11,22^. *ΔhtkB* cells were grown for 24 h under the same conditions as KU216 to stationary phase (Supp. Fig. 2), followed by cryo-EM grid preparation, data acquisition, and cryo-ET processing. Visual inspection of tomograms suggested that stationary-phase *ΔhtkB* chromatin contained smaller hypernucleosomes than the parental KU216 strain (Fig. 2a, Video S2). STA identified six classes of hypernucleosomes, corresponding to particles wrapping ~60-210 bp of DNA (Fig. 2b, c, Supp. Fig. 3), revealing a strong shift toward shorter particles relative to those in stationary-phase KU216. Mapping the subtomogram averages back into the tomograms based on their positions and orientations further confirmed that *ΔhtkB* chromatin fibers are composed predominantly of smaller hypernucleosomes than those of the parental strain (Fig. 2a, d).

**Fig. 2.**
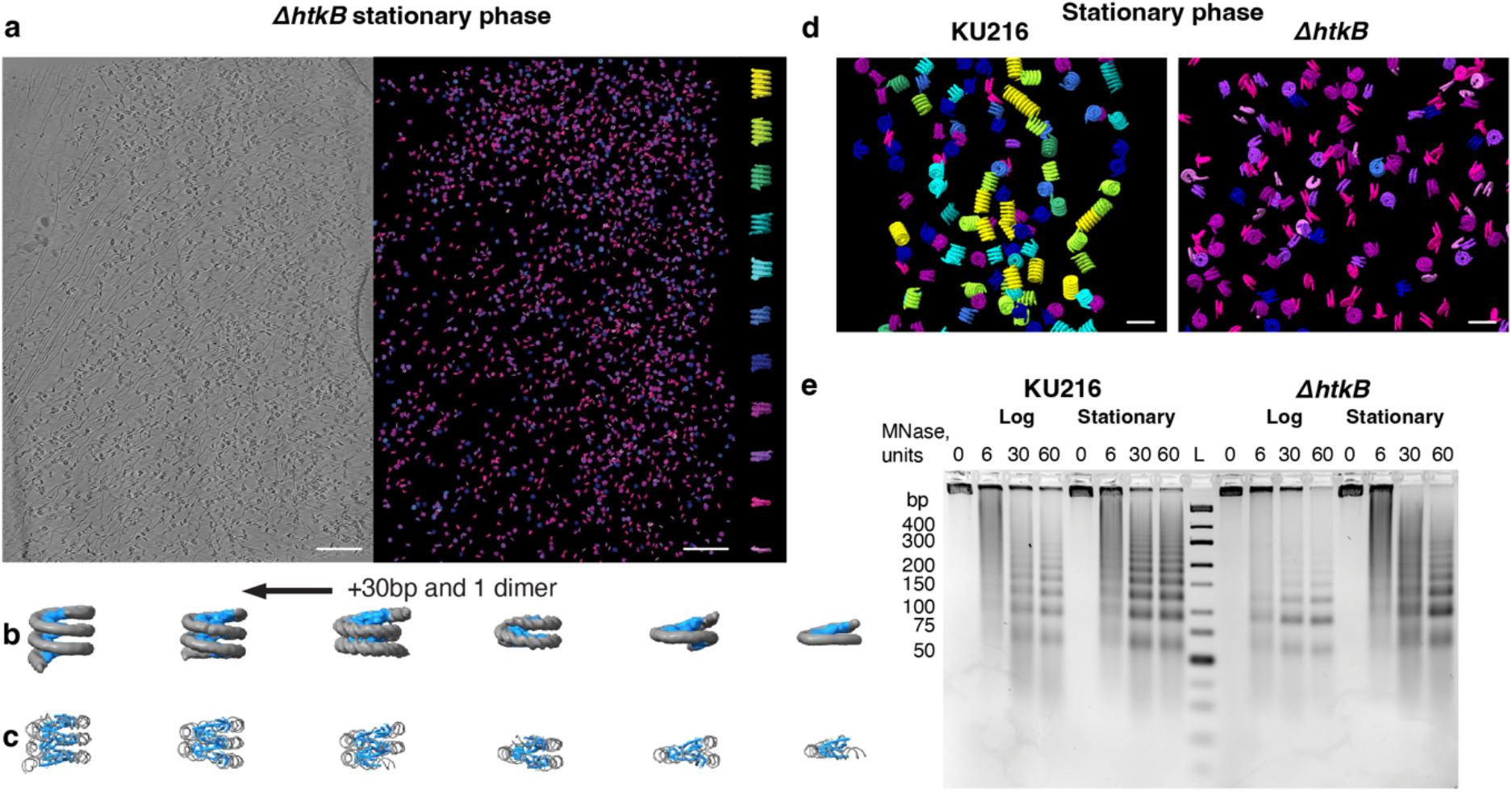
Cryo-ET of *T. kodakarensis* chromatin from stationary-phase *ΔhtkB* cells. a. Representative slice through a cryo-ET tomogram. Left: raw tomographic slice; right: subtomograms pasted back into the tomogram at their refined positions and orientations. Scale bar - 100 nm b. Six STA maps at resolutions of 8.2-15.4 Å, with DNA shown in grey and histones in blue. c. Corresponding structural models of the different hypernucleosomal classes, with DNA shown in grey and histones in blue. d. MNase-digest of chromatin: from left to right: KU216 cells in log-phase, KU216 cells in stationary phase, *ΔhtkB* cells in log-phase, *ΔhtkB* cells in stationary phase. For each sample, four lanes are shown with increasing amounts of MNase, ladder lane is marked with L.

To validate these findings using an orthogonal approach, we performed MNase digestion on chromatin from stationary-phase KU216 and *ΔhtkB* cells (Fig. 2e). In KU216, MNase digestion produced the characteristic ladder of protected DNA fragments ranging from ~60 bp to >400 bp, with a periodicity of ~30 bp. In contrast, the *ΔhtkB* sample showed a clear enrichment of shorter fragments, with the strongest bands at ~90–120 bp, and the ladder did not extend beyond ~300 bp, consistent with the cryo-ET data. Although our initial analysis focused on stationary phase, we also included log-phase KU216 and *ΔhtkB* samples in the MNase digest experiments, and observed the same overall trend: KU216 chromatin displayed a broader range of protected DNA fragments, whereas *ΔhtkB* chromatin was enriched in shorter species (Fig. 2e). Together, these results show that absence of HTkB shifts chromatin toward substantially shorter hypernucleosomes in both log and stationary phases.

### *T. kodakarensis* chromatin geometric analysis reveals local chromatin organization

Subtomogram averaging provides not only structures of individual hypernucleosomes, but also the opportunity to analyze their relative positions and orientations within tomograms. We therefore carried out geometric analysis on stationary-phase KU216 and stationary-phase *ΔhtkB* chromatin. This analysis used 5 tomograms from the KU216 dataset and 3 tomograms from the *ΔhtkB* dataset, with comparable numbers of particles (Methods, Table 1). To maximize particle identification completeness and accuracy, template-based particle picking was followed by manual curation of each tomogram prior to further analysis.

**Table 1.**
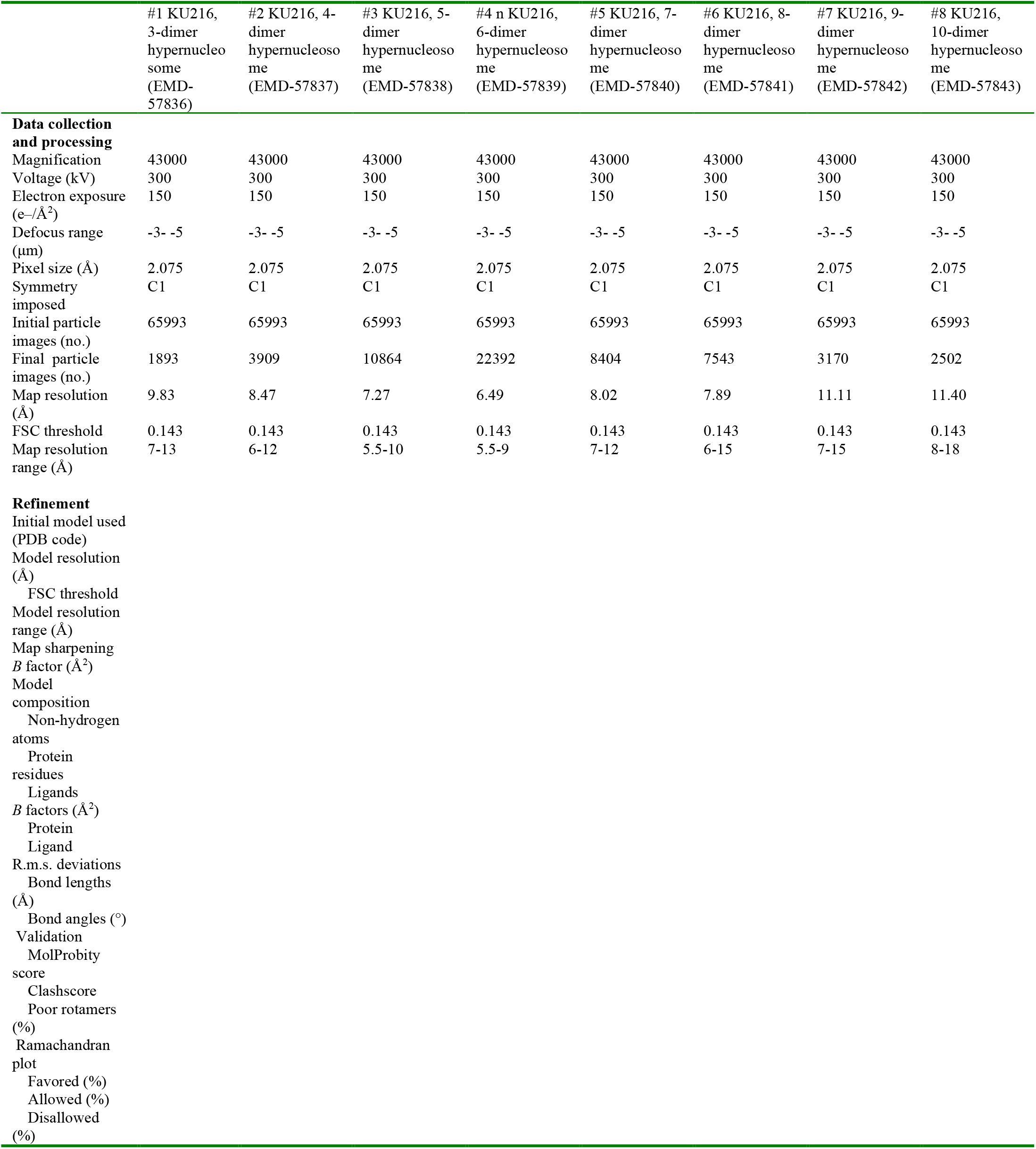
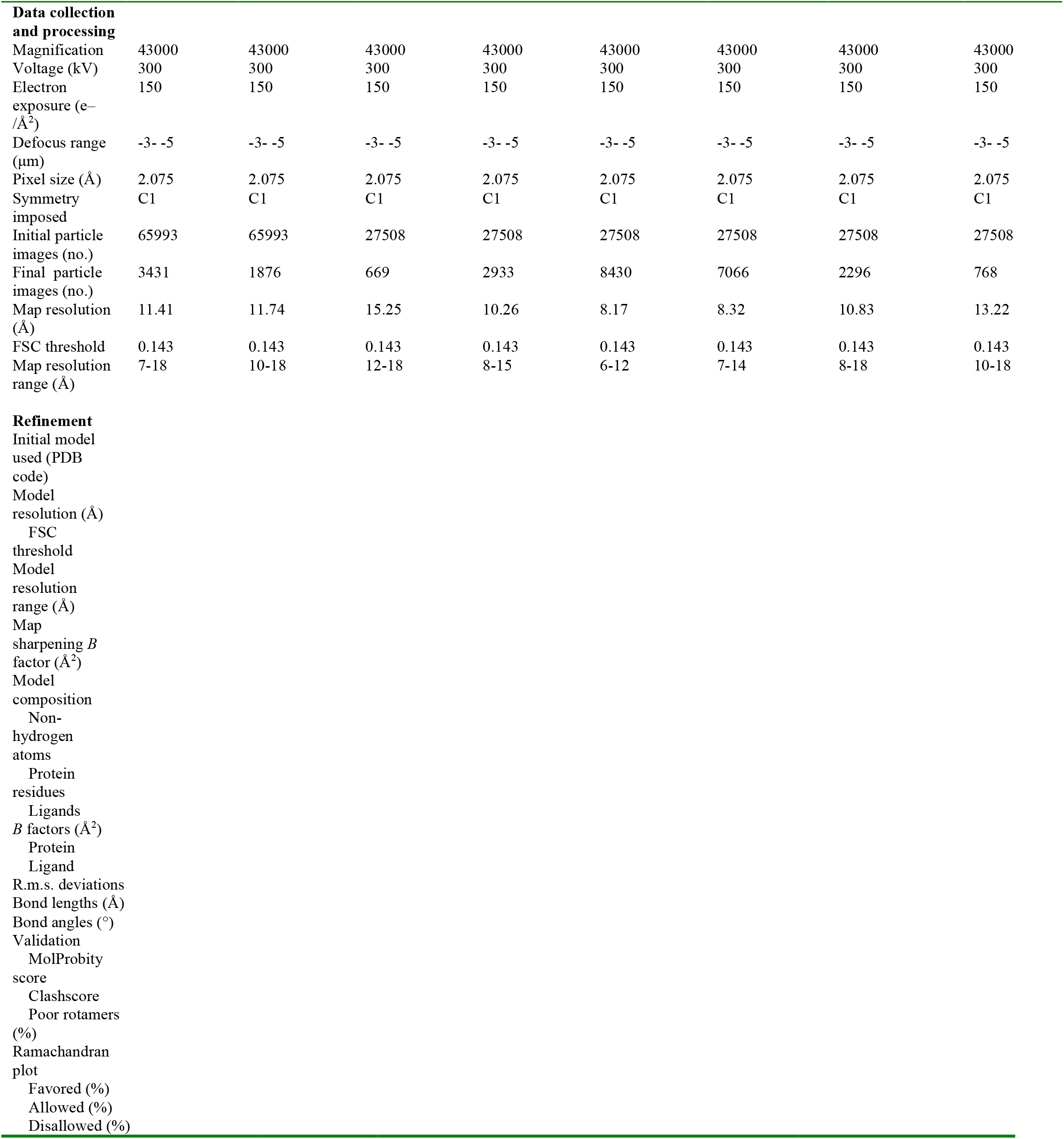
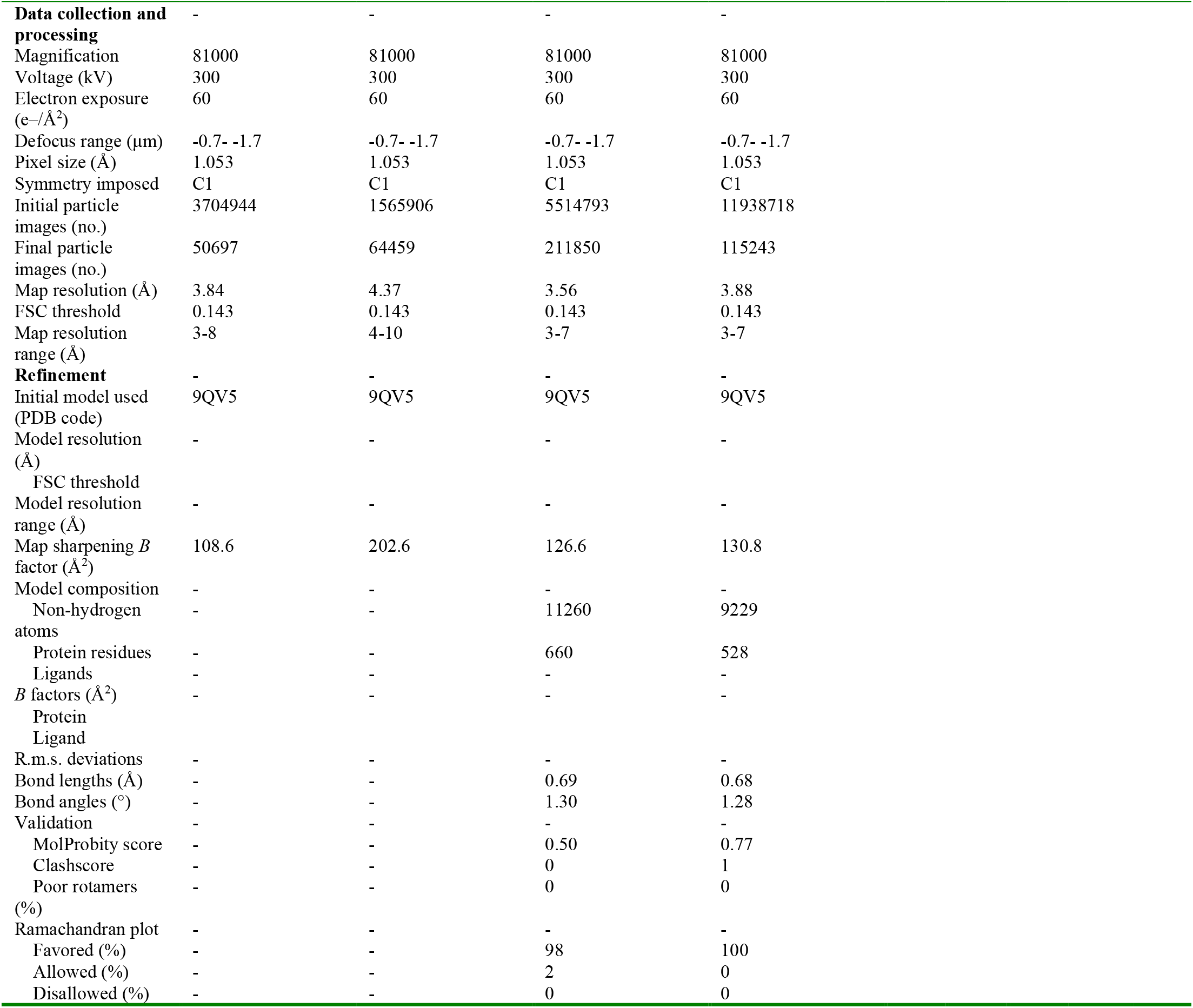
Cryo-EM data collection, refinement and validation statistics.

We first calculated pairwise distances between hypernucleosomal particles. In both datasets, this revealed a clear peak at approximately 12 nm (Fig. 3a), distinct from simulated random particle distributions with the same particle numbers per tomogram. In KU216, this peak was sharp, whereas in *ΔhtkB* it was broader, indicating reduced local order in the absence of HTkB histone.

**Fig. 3.**
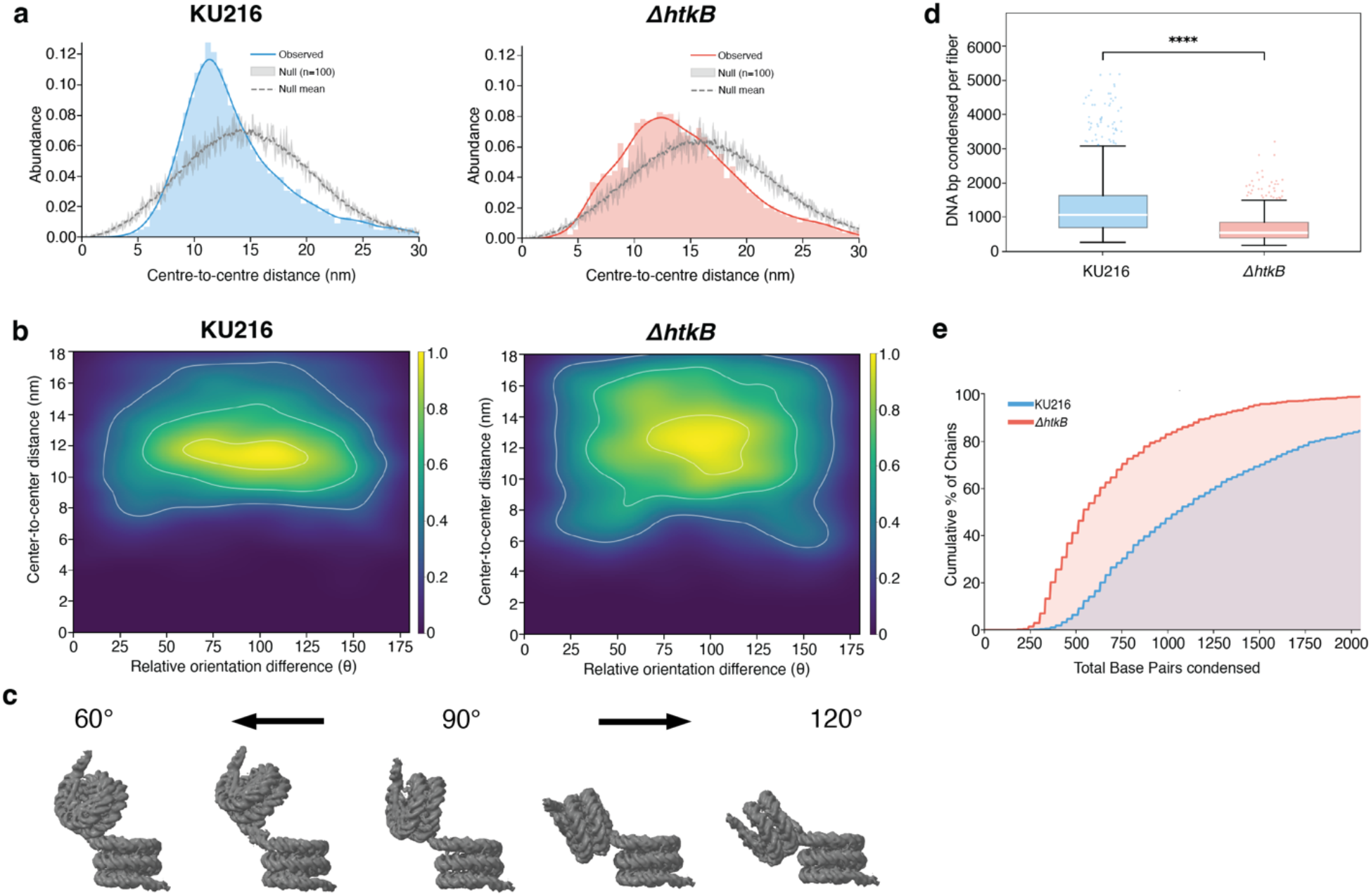
Geometric analysis of local chromatin organisation based on cryo-ET data. a. Pairwise distance distributions of neighbouring hypernucleosomal particles, measured between centers of particles. b. Pairwise relative orientation distributions of neighbouring hypernucleosomal particles. c. Representative examples of different local orientations, illustrating the continuum of slinky configurations from acute to obtuse angles. d. Chromatin fiber content analysis: total DNA length in bp, condensed into a single fiber. e. Percentage of fibers that have a certain length of DNA (bp) condensed.

We next examined the relative orientations of neighboring hypernucleosomes to define the nature of local contacts within archaeal chromatin fibers. This analysis revealed strong enrichment for particle pairs oriented at angles of ~60–120° relative to one another (Fig. 3b). However, rather than adopting a single defined geometry, such as the previously described ~90° slinky arrangement^16^, neighboring hypernucleosomes formed a continuum of relative orientations spanning acute to obtuse angles in the slinky-like configurations^16^ (Fig. 3b, c). Overall, local contacts extended across a broad angular range, from ~20° to ~160° and beyond (Fig. 3b). In *ΔhtkB*, the overall distribution of relative angles was similarly broad, and the main peak was likewise centered at ~60-120°, but the distribution was more diffuse and the corresponding pairwise distance distribution was broader, consistent with less ordered contacts between neighboring hypernucleosomes in the absence of HTkB.

Finally, we grouped neighboring hypernucleosomes separated by minimal distances to identify particles belonging to the same chromatin fiber. This allowed us to determine the number of hypernucleosomes within individual fibers and to estimate the total amount of DNA wrapped per fiber. Although the number of particles per fiber was only modestly reduced in *ΔhtkB* (Supp. Fig. 4), the total amount of wrapped DNA differed substantially between the two strains (Fig. 3d, e). In KU216, 50% of fibers contained at least ~1000 bp of wrapped DNA, whereas in *ΔhtkB* the corresponding value was only ~500 bp. Thus, although *ΔhtkB* fibers often contain similar numbers of particles, they are composed of shorter hypernucleosomes and therefore package substantially less DNA overall.

### 2D cryo-EM reveals condition-dependent shifts in hypernucleosome size distributions

Cryo-ET and STA provide molecular identities and enable discrimination between nucleosome species across a broad size range, but this approach is time-consuming and yields a comparatively limited number of particles. MNase digestion, on the other hand, provides a rapid readout of protected DNA fragments, but has important limitations. Fragments larger than ~400 bp are poorly resolved, and, in principle, protected DNA fragments may arise not only from histone-bound species but also from protection by other DNA-binding proteins. Therefore, to complement these methods, we collected 2D cryo-EM datasets on chromatin isolated from stationary-phase KU216 and stationary-phase *ΔhtkB* cells. Following particle picking, 2D classification was performed (Fig. 4a). Side-view classes were grouped according to hypernucleosome size, and the number of particles contributing to each class was quantified (see Methods, and Supp. Fig. 5). This analysis revealed a clear effect of *htkB* deletion, with a pronounced shift toward shorter hypernucleosomal particles (Fig. 4b). Whereas ~57% of hypernucleosomes in stationary KU216 contained six, seven, or more histone dimers, only a small fraction of 6-7-dimer particles was present in the stationary *ΔhtkB* dataset (~16%), and larger particles were not detected (Fig. 4a). These results are consistent with the distributions observed by both cryo-ET and MNase digestion, validating this as an orthogonal approach for assessing hypernucleosome size distributions.

**Fig. 4.**
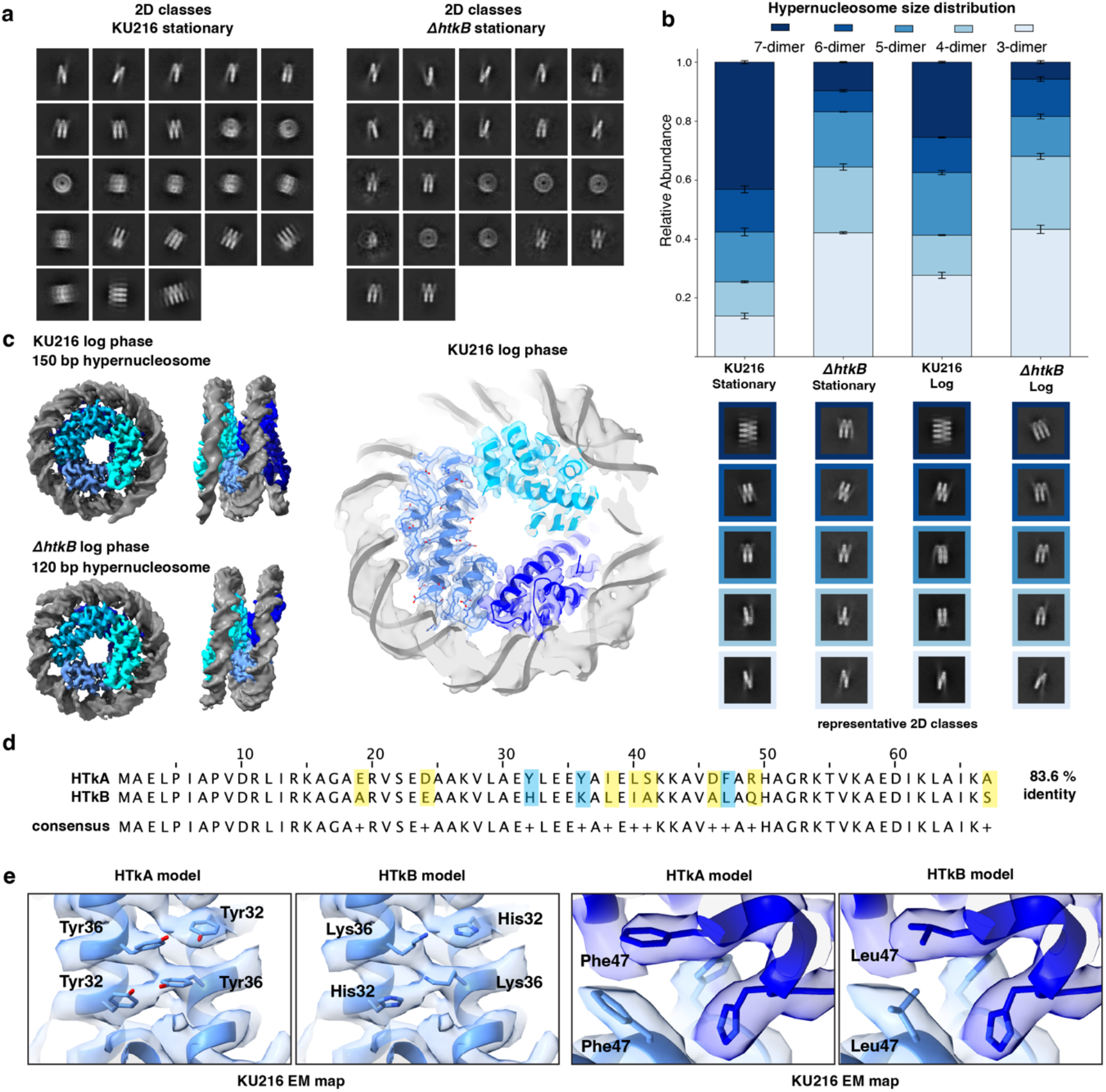
Single particle analysis and quantification. a. 2D classes from KU216 stationary and *ΔhtkB* stationary cryo-EM SPA datasets. b. Quantification of hypernucleosome sizes in different datasets. Below – representative 2D classes. c. The most abundant (highest resolution) class of particles is shown for each of the two datasets: 5-dimer 150 bp for the log-phase KU216 dataset and 4-dimer 120bp for the KU216 *ΔhtkB* dataset. On the right – a close up of 150 bp EM map with the model fitted. d. Sequence alignment of HTkA and HTkB histone sequences. Consensus alignment is shown below, residues that are different are highlighted with colour (yellow). Residues that are visualised below in panel e are highlighted in blue. e. Close up of the KU216 log phase EM map with either the HtkA or HTkB model fitted.

We next applied the same workflow to chromatin isolated from log-phase KU216 and log-phase *ΔhtkB* cells, completing the set of cryo-EM data for both growth phases and both genetic backgrounds (Supp. Fig. 5). Comparison of these datasets showed that chromatin from log-phase KU216 had shorter hypernucleosomes relative to stationary-phase KU216 (Fig. 4b). For example, only ~36% of hypernucleosomes in log-phase KU216 contained more than six histone dimers, corresponding to >180 bp of wrapped DNA, whereas this proportion was ~57% in stationary phase. In contrast, the difference between log-phase and stationary-phase *ΔhtkB* was less pronounced, with both datasets enriched in shorter hypernucleosomal species, showing a strong effect of *htkB* deletion. Thus, both growth phase and absence of HTkB histone influence hypernucleosome size distributions, with the strongest enrichment of longer hypernucleosomes observed in stationary-phase KU216, and the shortest in log-phase *ΔhtkB*.

### SPA reconstruction reveals high-resolution structures of native hypernucleosomes

The four cryo-EM datasets (KU216 and *ΔhtkB*, in stationary and log-phases) were subsequently processed using a complete SPA workflow (Methods, Supp. Fig. 6-9). Following heterogeneous refinement, 3D classifications and non-uniform refinement, multiple medium- to high-resolution maps were obtained for hypernucleosomes of different sizes (Table 1, Supp. Fig. 6-9). In all datasets, the resulting SPA reconstructions were consistent with the structures observed by cryo-ET but were better resolved (Supp. Fig. 10). Notably, hypernucleosomes in all SPA maps adopted closed conformations (Supp. Fig. 6-9). A small degree of breathing was observed, manifested as a slight increase in spacing between adjacent DNA gyres (Video S3, S4), but this flexibility was limited, and no open conformations comparable to those reported for other archaeal species were detected^17^.

The log-phase datasets contained more particles overall and therefore yielded the highest-resolution reconstructions. In the KU216 log-phase dataset, the most abundant class corresponded to a 5-dimer hypernucleosome resolved at 3.56 Å (Fig. 4c, Supp. Fig. 10). In the *ΔhtkB* dataset, the most abundant class corresponded to a 4-dimer hypernucleosome resolved at a slightly lower resolution of 3.88 Å (Fig. 4c, Supp. Fig. 10). Both reconstructions showed clear side-chain densities. At this resolution, sequence-dependent differences between HTkA and HTkB could also be assessed. In the KU216 dataset, densities reflect the averaged contribution of both histones, whereas the *ΔhtkB* dataset contains only HTkA. Although HTkA and HTkB are 83.6% identical, several residues differ between them (Fig. 4d). For example, HTkA contains Tyr32 and Tyr36, where HTkB contains His32 and Lys36, the density in the map of KU216 remains ambiguous even at high resolutions, because the KU216 parent strain contains a mixture of HTkA and HTkB proteins, that get averaged out (Fig. 4d, e). These data show that SPA of chromatin isolated directly from archaeal cells enables high-resolution structural analysis of native hypernucleosomes without in vitro reconstitution.

Together, these results show that archaeal chromatin architecture varies with both growth phase and histone composition, revealing a dynamic and condition-dependent organization of the genome. Overall, we show visualize near-native archaeal chromatin at high resolution across different cellular states, and demonstrate that archaeal chromatin architecture changes in response to growth phase and the presence or absence of histone HTkB, revealing a dynamic, condition-dependent organization of the archaeal genome.

## Discussion

This study provides the first high-resolution structural analysis of chromatin isolated directly from archaeal cells. Using *T. kodakarensis* as a model system, we show that chromatin adopts a variable-beads-on-a-string architecture in which each bead corresponds to a closed hypernucleosome of variable length. This general model was previously proposed based on MNase digestion patterns and supported by an X-ray structure of the minimal nucleosome, showing that an archaeal histone dimer binds ~30 bp of DNA^1,2^. However, those approaches had important limitations: MNase digestion does not identify the protein responsible for DNA protection, whereas previous X-ray and EM studies relied on reconstituted systems with artificial DNA substrates and typically examined one type of particle at a time^1,16^.

Here, we address these limitations by analysing chromatin isolated directly from archaeal cells. Our ex situ approach preserves native chromatin composition while avoiding biases associated with in vitro reconstitution, including artificial DNA sequences and non-physiological histone stoichiometries. By combining cryo-EM and cryo-ET, we obtain both high-resolution information on chromatin building blocks and geometric information on local fiber organization. Together, these data identify hypernucleosomes as fundamental structural units of archaeal chromatin and support a variable-beads-on-a-string model for chromatin fiber organization (Fig. 5). Similar ex situ strategies have proven informative for eukaryotic chromatin^23^, where structures and local contacts identified outside the cell were subsequently observed in situ^24^. In the future, this approach can also be applied to study chromatin of other archaeal species in a straight-forward manner, without the need for additional sample preparation steps or extra equipment, like focused-ion-beam milling, and can overcome the challenges of poor intracellular contrast that is well known for many archaeal species^25^.

Our data further show that hypernucleosomes span a broad size range across all datasets, from 60 bp to 360 bp of wrapped DNA, yet consistently adopt closed conformations characterized by tightly packed DNA gyres and strong stacking interactions. This finding is consistent with previous in vitro work^16^ and predictions^26^ and supports the view that compact, closed hypernucleosomes represent the predominant architecture in *T. kodakarensis* and potentially in many other archaea. Importantly, we do not detect the open conformation recently reported for the Asgard histone HHoB^17^, indicating that this open state is not a general feature of archaeal chromatin and instead reflects lineage-specific properties. At the same time, we observe limited “breathing” of hypernucleosomes within our datasets, visible as small changes in spacing between adjacent DNA gyres. Such behaviour is characteristic of nucleosome dynamics in both archaeal and eukaryotic systems^16,27^, but here it is observed within continuous chromatin fibers rather than isolated particles. Thus, archaeal chromatin appears to combine substantial compaction with a small degree of local conformational flexibility.

Having established the basic structural features of archaeal chromatin in stationary-phase *T. kodakarensis*, we next examined the role of histone paralog composition using an *htkB* deletion mutant. A regulatory function for archaeal histone paralogs has long been proposed in species encoding multiple histones^28,29^. Biophysical work has shown that HTkB has slightly higher affinity to DNA compared to HTkA^30^, and analogous histone pairs in other archaea have also been reported to differ in the rigidity of the chromatin fibers they form^12^. In addition, the relative abundance of histone paralogs can vary with growth phase (i.e. in *M. fervidus*)^31^, suggesting that paralog composition may contribute to chromatin regulation in vivo. However, a direct connection between histone paralog identity, cellular state, and chromatin architecture has remained unclear.

Our data now provides such a link. In both log and stationary phases, the *ΔhtkB* mutant is shifted toward substantially shorter hypernucleosomes than the parental strain, with the difference becoming especially pronounced in stationary phase. These results indicate that HTkB promotes the formation or stabilization of longer hypernucleosomes in vivo. In its absence, cells fail to accumulate the long hypernucleosomal species that are prominent in stationary-phase wild type. This functional role is consistent with previous evidence that deletion of HtkB mildly slows growth in *T. kodakarensis*^*11*^, and supports the idea that histone paralogs are not functionally redundant but instead make distinct contributions to chromosome organization.

Beyond the structures of individual particles, our analysis also provides insight into local chromatin fiber organization. Although long-range chromatin arrangement is disrupted in crude lysates, short-range nucleosome-nucleosome contacts remain preserved. We observe enrichment of closely apposed hypernucleosomes and a continuum of relative orientations between neighboring particles rather than a single dominant arrangement. This suggests that local fiber geometry is flexible. Notably, the *ΔhtkB* mutant shows a broader distribution of relative angles, indicating reduced local order. One possible explanation is that the increased abundance of shorter hypernucleosomes in this background reduces steric constraints, thereby permitting a wider range of local configurations. Together, these observations suggest that HTkB contributes not only to hypernucleosome length but also to the local organization of chromatin fibers.

Another major conclusion from this work is that archaeal chromatin architecture changes with growth phase. In the parental strain, stationary-phase cells are enriched in longer hypernucleosomes relative to log-phase cells, indicating that chromatin becomes more compact as growth slows. This provides a structural basis for earlier suggestions that archaeal chromatin organization changes with physiological state^19^. It is also consistent with previous observations that transcription is strongly reduced in stationary-phase *T. kodakarensis*^*32*^ and that chromatin appears more compact by low-resolution AFM imaging^19^. The data support a model in which increased hypernucleosome length may contribute to a more repressive, condensed chromatin state during stationary phase.

More broadly, our findings suggest that modulation of hypernucleosome length may represent a central mechanism by which archaea regulate DNA accessibility. In this model (Fig. 5), smaller hypernucleosomes predominate during active growth, when replication and transcription are high, whereas longer hypernucleosomes accumulate under nutrient limitation and reduced proliferation, promoting chromatin compaction. HTkB appears to play an important role in this transition by enabling the formation of longer, more ordered chromatin assemblies.

**Fig. 5.**
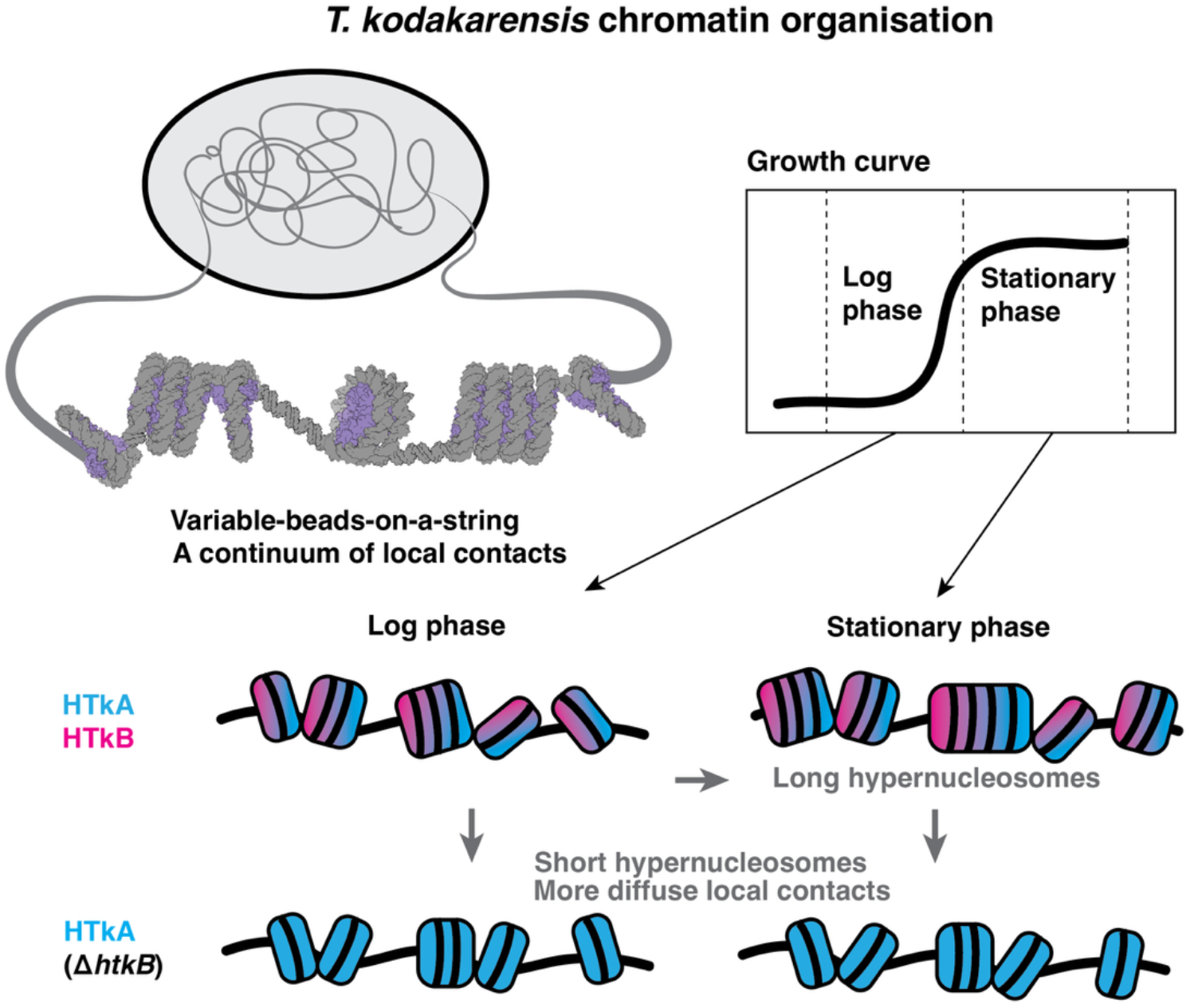
Model of *T. kodakarensis* chromatin organisation. *T. kodakarensis* chromatin forms a variable-beads-on-a-string fibers with a continuum of local contacts. In stationary phase, HTkA/HTkB-containing chromatin consists of longer hypernucleosomes, whereas log-phase chromatin fibers contain shorter hypernucleosomes. When only HTkA histone is present (*ΔhtkB*) - the hypernucleosomes become shorter in both log and stationary phases, and the local chromatin contacts - more diffuse.

In summary, we show at high resolution the organization of archaeal chromatin directly isolated from cells, and show how chromatin architecture changes across growth conditions and histone composition. These findings establish a framework for understanding archaeal genome organization at molecular resolution and provide direct support for a variable-beads-on-a-string model built from closed hypernucleosomes of variable size. Future work combining ex situ structural analysis with in situ cryo-ET and orthogonal genomics approaches should help bridge local chromatin structure with chromosome-wide organization and function.

## Acknowledgments

This work is supported by the European Union (ERC Starting Grant 3DchromArchaea, grant agreement no. 101076671, to S.O.D). H.A. was supported by JSPS KAKENHI grant number JP25H00432.

We thank: Joseph Bartho for technical assistance with microscopy; Thomas Hoffmann and EMBL IT for IT support; Higor Rosa, Euan Pyle and Linhua Tai for helpful discussions and sharing scripts related to tomography processing; Katharina Werner for help with the EPU Multigrid setup.

Funded by the European Union. Views and opinions expressed are however those of the authors only and do not necessarily reflect those of the European Union or the European Research Council Executive Agency. Neither the European Union nor the granting authority can be held responsible for them.

## Author contributions

M.D and F.R cultivated the archaeal cells with help from T.S.B. M.D. prepared cryo-EM grids, collected single particle and tomography data with help of Q.D and S.U and processed data with input from T.S.B, Q.D and S.O.D. M.D built models with support from Q.D. MNase digest were performed by F.R with input from N.T. S.Y. generated the HTkB deletion mutant strain under supervision of N.T and H.A. S.O.D conceived the study and supervised work. M.D. and S.O.D. wrote the manuscript with input from all authors.

## Data availability

All EM maps from STA are deposited under EMDB IDs: EMD-57836 – 57853; Two highest resolution SPA EM maps are deposited as EMD-57854 and EMD-57855 with corresponding molecular models PDB IDs 30KF and 30KG.

