## Supplementary Files for "Cryo-electron tomography reveals near-native archaeal chromatin architecture"

**Supplementary Information contains:**

- Materials and Methods
- Supplementary Figures S1-S10

#### Materials and Methods

##### Cultivation of *Thermococcus kodakarensis*

*T. kodakarensis* KU216 was cultured in 20 mL glass bottles as previously described[33,34]. Briefly, *T. kodakarensis* was kept under anaerobic conditions at a temperature of 85ºC in rich growth media supplemented with 2 g/L elemental sulfur and 0.8 mg/L Resazurin (ASW-YT-Sº) (Artificial Sea Water – Yeast extract (Sigma Aldrich) + Tryptone (Oxoid) – elemental sulfur). 5 µL of 0.66 g/mL $Na_2S•9H_2O$ was added to the media until it appeared light yellow. Fresh, preheated media were inoculated at an $OD_{600}$ of 0.01 and grown for either 5 h, when cells were harvested during exponential growth (log-phase), or 24 h, when cells were harvested in stationary phase.

##### Construction of *T. kodakarensis htkB* deletion strain

Marker-less deletion of *htkB* (*TK2289*) in *T. kodakarensis* was performed using a pop-in/pop-out approach[34]. *T. kodakarensis* KU216 was used as a parental strain, lacking the orotidine-5'-monophosphate decarboxylase gene *pyrF*. The Δ*pyrF* mutation confers uracil auxotrophy as well as resistance to 5-fluoroorotic acid (5-FOA). To construct a deletion plasmid targeting *htkB*, ~1-kb regions immediately upstream and downstream of the coding sequence were PCR-amplified using genomic DNA from KU216 as a template. These fragments were cloned into the SphI-BamHI site of the pUD3 vector containing the *pyrF*[35] marker using In-Fusion Snap Assembly Master Mix reagents (Takara Clontech). The coding sequence of *htkB* was then removed from the resulting plasmid by inverse PCR, and the PCR product was circularized using In-Fusion Snap Assembly Master Mix reagents to generate the *htkB* deletion plasmid). The resulting deletion plasmid was used to transform KU216, allowing integration of the construct into the target genomic locus. Transformants with the *pyrF* marker were selected in the uracil-free medium ASW-AA-m1-S0[36].To select strains in which *pyrF* had been lost via pop-out, cells were further grown in solid ASW-YT-m1-S0 medium[37] supplemented with 8.3 g/L of 5-FOA•H2O and 44 mM NaOH. Colonies resistant to 5-FOA were analyzed by PCR to confirm marker-less deletion of *htkB*.

##### Micrococcal Nuclease Digestion

MNase (micrococcal nuclease) digestion was performed as previously described with slight modifications[2]. Briefly, 200–400 ml of *T. kodakarensis* culture were placed on ice and then harvested by centrifugation. Next, cells were resuspended in 600 µL 0.8× ASW and stored at −20 °C until further processing. Frozen samples were thawed on ice and 1350 µL extraction buffer (25 mM HEPES pH 7.0, 15 mM $MgCl_2$, 100 mM NaCl, 0.4 M sorbitol, 0.5% Triton X-100 (Merck)) was added. Samples were incubated on ice for 10 min, then centrifuged at 14,000 × g for 20 min at 4 °C. The supernatant was discarded and the chromatin-containing pellet was resuspended in 1×MNase buffer (New England Biolabs NEB; 50 mM Tris-HCl, 5 mM $CaCl_2$, pH 7.9).

MNase digestion reactions were performed in a final volume of 200 µL containing 0, 6, 30, or 60 units of MNase (NEB). Reactions were incubated for 30 min at 37 °C with shaking at 1000 rpm. Digestion was stopped by adding EDTA to a final concentration of 25 mM. Samples were then supplemented with SDS to a final concentration of 0.8% (Sigma-Aldrich) and proteinase K to a final concentration of 0.5 mg/mL (NEB) and incubated for 3 h at 37 °C.

DNA was extracted using phenol:chloroform:isoamyl alcohol (25:24:1; Carl Roth) and precipitated with 100% ethanol and 0.3 M sodium acetate (Carl Roth). After 1h at −20 °C, DNA was pelleted by centrifugation at 20,400 × g for 30 min at 4 °C. The supernatant was removed, and the pellet was washed with 1 mL 70% ethanol. DNA was dissolved in 10 mM Tris-HCl (pH 7.5) containing 0.05 mg/mL RNase A (Thermo Fisher Scientific). DNA concentration was determined using the Qubit dsDNA High Sensitivity Assay Kit (Thermo Fisher Scientific). 2 µg DNA per sample was separated on a 4% NuSieve GTG agarose gel (Lonza) in 1× TBE buffer at 100 V for 90 min. DNA ruler TriDye Ultra Low Range DNA Ladder (NEB) was used. DNA fragment distribution was quantified using Fiji software[38].

##### Cryo-EM Sample Preparation

*T. kodakarensis* KU216 or *T. kodakarensis* Δ*htkB* cells were harvested at 10,000 g for 5 min. To gently lyse cells, 0.4x ASW supplemented with 10 µg/mL RNAse A (Thermo Fisher Scientific), and 0.05% Triton-X100 (Merck) was added to the cell pellet, and cells were gently resuspended. Cell lysate was kept at the cultivation temperature until further sample processing (maximally for 20 minutes).
Next, Quantifoil R3.5/1 CU 300 mesh or UltrAuFoil R2/2 Au 200 mesh grids were glow-discharged for 45 seconds from both sides at 25 mA using a GloCube Plus (Quorum Technologies). Samples were plunge-frozen using a Vitrobot Mark IV (Thermo Fisher Scientific). 3.5 µL of lysed cell sample was applied to the front side of the cryo-EM grids. The Vitrobot chamber was set to 20 ºC at 100% humidity. The sample was blotted from the back side between 7.5-15 seconds with a blot force of 20 and a wait time of 3 seconds. A custom Teflon sheet was used to avoid frontside blotting.

**Cryo-electron Tomography Data Collection and Processing**

Tilt-series of *T. kodakarensis* cell lysates were collected on Titan Krios G3 microscope operated at 300 keV equipped with a field emission gun, K3 direct electron detector (Gatan Inc), and a Bioquantum energy filter. Data were collected at a nominal magnification of 42,000x corresponding to a pixel size of 2.075 Å/px with defocus ranging between -3 and -5 µm following a dose-symmetric tilt-scheme[39], with 3° increments between 51° and +51° stage tilt and a total dose of 150 $e^-$/Å2. All data was processed in WarpTools[40,41] (https://warpem.github.io/) if not indicated otherwise. Motion correction and CTF estimation were performed with default parameters suggested in WarpTools documentation. Initial tilt stacks were manually corrected for bad tilts using a custom script (kindly provided by Linhua Tai), followed by alignment using the WarpTools implementation of AreTomo[42] (https://github.com/czimaginginstitute/AreTomo3). Tomograms were reconstructed at a pixel size of 10 Å and used for particle localization and subsequent subtomogram averaging. For visualization, tomograms were denoised using cryoCARE[43] (https://github.com/juglab/cryoCARE_pip).

**Subtomogram averaging**

For the *T. kodakarensis* KU216 stationary-phase dataset, for initial reference generation from the data, a cylinder was created in IMOD[44,45] and used for template matching in PyTom[46] (https://github.com/SBC-Utrecht/pytom-match-pick) at 10 Å/px. Overall workflow is shown in Supp. Fig. 1. Subtomograms were generated in WarpTools at a pixel size of 10 Å/px and imported into Relion 4.1[47]. Two rounds of 3D-classification to remove false positive particles identified during template matching were followed by a 3D-refinement. The resulting subtomogram average was then used to run the next round of template matching in PyTom. Particles were again extracted at a pixel size of 10 Å/px and imported into Relion4.1. An initial 3D-classification was performed to remove false-positive picks, followed by classification to resolve distinct hypernucleosome sizes, yielding five different classes. Particles of each class were re-extracted at 5 Å/px and refined again. Particles were then unbinned to their original pixel size of 2.075 Å/px and used as input for M[48]. Initially, only image shifts and deformations in the tilt series were corrected to avoid overfitting parameters. After M, the CTF was re-estimated, and tomograms were reconstructed again with optimized parameters from M. Template matching was performed with most abundant subtomogram average from the previous iteration at a pixel size of 10 Å/px. Subtomograms were re-extracted and used for subsequent subtomogram averaging. Initial 3D-classification yielded clean classes of hypernucleosomes of several sizes, and all particles from template matching were used for subtomogram averaging. Hypernucleosome particles were separated by size, with a separation of 0.5 turns corresponding to 1 additional histone dimer with 30 additional bp per class. In total, eight different classes were identified and further refined. After re-extraction and 3D-refinement at 5 Å/px, particles were unbinned and imported into M for multi-species refinement. After the second refinement in M, PyTom template matching was re-run with a wider mask to include as many nucleosomal particles as possible. Particles were re-classified as described above. Particles of each class were then re-extracted at a pixel size of 5 Å/px and 3D-refined. To remove duplicates between different hypernucleosome classes, particle sets were compared with a custom Python script and overlapping particles were removed hierarchically from larger to smaller particles. This process was repeated until no duplicates remained across any particle lists. Additional picks were added manually for a limited number of tomograms in ArtiaX[49] used for the geometrical analysis to ensure the most complete particle coverage. To ensure manually picked particles were assigned to the right class, similarly sized particles were pooled, and a 3D-classification of each pooled group was performed, and misassigned particles were re-ordered (Supp. Fig. 1). This yielded 10 distinct nucleosome sizes wrapping between 90-360 bp of DNA per hypernucleosome. All particles were unbinned again and used for two rounds of multi-species refinement in M, yielding subtomogram averages up to 6.5 Å resolution.

For the *T. kodakarensis ΔhtkB* stationary-phase, a similar approach as described above was followed. (Supp Fig. 3). Instead of an initial reference generation by template matching with a cylinder, the initial 5-dimer hypernucleosome subtomogram average of the *T. kodakarensis* KU216 stationary-phase dataset was used for the first iteration of template matching in PyTom, after two rounds of 3D-classification and an initial 3D-refinement, three different hypernucleosome sizes were re-extracted at 5 Å/px and 3D-refined, before being imported into M at a pixel size 3 Å/px of for correction of image shifts and deformations in the tilt series. Afterward, template matching was performed again in PyTom on the corrected tomograms. Extracted particles were 3D-classified and refined twice, followed by re-extraction and another refinement at 5 Å/px. For a second M refinement, the resulting four different hypernucleosomes were used. Following M refinement template matching in PyTom was re-run, this time with a wide mask to make the particles cutoff more permissive. After two rounds of 3D classification, particles were merged with manual picks (ArtiaX) from three tomograms used for the geometrical analysis. Re-classification of all particles revealed two more hypernucleosome classes. All six classes were refined at 5 Å/px and imported into M at a pixel size of 3 Å/px. As for the KU216 dataset, two rounds of M refinement were performed, resulting in hypernucleosome reconstructions ranging from 8.17 Å to 15.25 Å, with 2-7 dimers wrapping 60-210 bp of DNA.

**Geometrical Analysis of Chromatin Fibers**

Particle picking and subtomogram averaging were performed as described above to obtain the spatial orientation of hypernucleosome particles in the tomograms. Five tomograms of the *T. kodakarensis* KU216 stationary-phase dataset, and three tomograms of *T. kodakarensis ΔhtkB* stationary-phase dataset were manually curated after template matching following the second iteration of M, to get the most complete annotation of hypernucleosomes for each tomogram. Thus, after manual curation for five or three tomograms, the KU216 stationary dataset yielded 8251 subtomograms, and the Δ*htkB* stationary-phase dataset yielded 7078 subtomograms. After manual picking, particles were assigned to 1 of 10 size classes (KU216 stationary phase) or 1 of 6 classes (KU216 *ΔhtkB* stationary-phase) (described above) via 3D classification.

For each particle class, a radius and height were assigned by assuming the hypernucleosomal particles could be represented as disk/cylinder particles. For this, the subtomogram average was aligned along the z-axis in ChimeraX[50,51] using the "measure inertia" command. After that, two particle endpoints were defined based on where the "first" and "last" histone dimers are located in the EM map. Distances were calculated from the center of mass to those points. This enabled reproducible calculation of hypernucleosome endpoints and measurement of distances between hypernucleosomes, without a bias towards particles with extended DNA densities. Start and end points were calculated for each hypernucleosome class and applied to each particle in the dataset.

***Pairwise distance calculation:*** For an initial assessment of particle distribution within the annotated tomograms, the nearest neighbors of each particle were calculated for both datasets (Fig. 3A). For this, subtomograms were extracted at a pixel size of 4 Å/px and refined in Relion. The resulting coordinates were then used to calculate the nearest neighbor within a 30 nm radius and compared to a simulated, random distribution of particles. Graphs for both datasets were generated with Python using matplotlib (3.10.5-gfbf-2025b).

***Pairwise relative orientation calculation:*** Based on pairwise distance calculations, a threshold of 18 nm was chosen to analyze the spatial relationships among particles. All particles were refined again at 4 Å/px in Relion using the same reference to ensure identical particle orientation across the different hypernucleosome classes in both datasets. To then understand relative orientation of the particles, their orientation difference θ (0-180°), the angle between disk normals $\hat{z}_i$ and $\hat{z}_j$, given by $\arccos(\hat{z}_i \cdot \hat{z}_j)$, was calculated (Supp. Fig. 4d). Graphs for both datasets were generated with Python using matplotlib (3.10.5-gfbf-2025b).

***Fiber length calculation**:* Chromatin fiber chains were identified by connecting spatially proximal nucleosomes based on surface-to-surface distance, which more accurately reflects physical contact than center-to-center distance given the cylinder-like hypernucleosome shape. Links between particles were accepted if the surface-to-surface distance fell under the threshold of 10 nm. Chains were then assembled by a greedy (exhaustive iterative) algorithm. If several surface-to-surface distances were under the 10 nm threshold, the smallest distance was chosen - this avoided branching of the chains within one fiber. Fiber length was calculated by summing the center-to-center distances of the particles contained within one chain. Total base pairs condensed per fiber were calculated by summing the per-class DNA content (60-360 bp for 2-12 dimer classes, assuming 30 bp DNA bound per nucleosome dimer). Graphs for both datasets were generated with Python using matplotlib (3.10.5-gfbf-2025b).

#### Cryo-EM Single Particle Analysis Data Collection and Processing

Data collection of log-phase KU216 and Δ*htkB* was performed on a Titan Krios G3 microscope operated at 300 keV equipped with a field emission gun, K3 direct electron detector (Gatan Inc) and a Bioquantum energy filter (Gatan Inc). Data were collected at a nominal magnification of 81,000x, corresponding to a pixel (px) size of 1.053 Å/px, with defocus ranging from -0.7 to -1.7 µm. The total electron dose of 60 e-/Å2 was distributed across 60 movie frames in electron-counting mode. The energy filter slit width was set to 20 eV.

Data collection for KU216 stationary-phase and Δ*htkB* stationary-phase was performed using the Multigrid implementation of EPU (Thermo Fisher Scientific). For each condition, 4500 micrographs were collected with the same parameters as mentioned above.

After collection, all data were imported into cryoSPARC 4.4.1[52]. After preprocessing and exposure curation, cryoSPARC's implementation of a blob picker with a particle size range of 80-300 Å was used to include a wide range of particles without bias. All particles were extracted at a pixel size of 2.1 Å/px with a box size of 128 px and subjected to several rounds of 2D classifications. If initial 2D classification revealed preferential picking of hypernucleosome side views, 5 classes of hypernucleosome top views from initial 2D classification were chosen as input for template picking or imported from another dataset if no initial top views were present in the 2D classifications. The template picker and blob picker outputs were merged, and duplicates were removed. Selected particles were subject to several rounds of ab-initio reconstruction, heterogeneous refinement, and non-uniform refinement (Supp Fig. 6-10). To generate final maps, the local resolution was estimated and a local filter applied, or maps were sharpened using a B-factor calculated from non-uniform refinement (Supp Fig. 6-10). Below are details for each data collection.

#### 2D Quantification of Hypernucleosome Sizes

Particles identified with the blob picker were cleaned using 2D-classification to remove false-positive picks. Next, for the quantification of hypernucleosome size, only side-view particles were selected. This allowed for consistent assignment of hypernucleosomal length without introducing template bias. The datasets were split into two equally sized parts, and side views were then re-classified into a distinct number of classes. The number of classes was determined by dividing the number of particles by 4000 (an arbitrary value shown to yield good classification results for quantification). (Supp. Fig 5). Classes were then assigned to five different hypernucleosome sizes based on their 2D average (3-dimer, 4-dimer, 5-dimer, 6-dimer and 7+-dimer – reflected in the number of visible hypernucleosome turns). Afterward, counts were converted to relative abundances by dividing each class by the total count across all size classes. Stacked bar charts were generated in Python using matplotlib (3.10.5-gfbf-2025b), with error bars representing the deviation between each split.

***KU216 stationary-phase dataset*:** After two 2D-classifications, 375,798 particles were reconstructed into four different volumes, followed by a heterogeneous refinement, resulting in three different-sized hypernucleosomes. Each class was subjected to a non-uniform refinement and re-extracted at the

original pixel size of 1.053 Å/px. After an initial non-uniform refinement, 3D-classification of the 6-dimer hypernucleosome and the 4-dimer hypernucleosome revealed two additional hypernucleosome classes (Supp. Fig. 6). The 8-dimer particles were reconstructed in three different volumes at a pixel size of 1.053 Å/px, followed by a non-uniform refinement of the high-resolution class of the heterogeneous refinement. Final non-uniform refinement, after reference-based motion correction, yielded five hypernucleosome classes with resolution up to 3.84 Å, with an initial consensus at 3.79 Å resolution for the 6-dimer and 3.88 Å for the 8-dimer (Supp. Fig. 6, 10). Final classes contained five different hypernucleosome structures, wrapping between 90 - 240 bp of DNA by 3-8 histone dimers.
***ΔhtkB Stationary-phase dataset*:** Data collected from Δ*htkB* stationary yielded 202,662 particles after two rounds of 2D-classification and were used for the reconstruction of five ab-initio classes (Supp Fig. 7). After a heterogeneous refinement, two hypernucleosome classes were refined and re-extracted at the original pixel size of 1.053 Å/px. The 4-dimer hypernucleosome class was subjected to another round of heterogeneous refinement, followed by a non-uniform refinement. After reference-based motion correction and a non-uniform refinement, the hypernucleosome classes for 3-dimer and 4-dimer reached resolutions of 4.37 Å and 6.3 Å, respectively, wrapping between 90 - 120 bp of DNA by 3-4 histone dimers. (Supp Fig 7)
***KU216 log-phase dataset*:** For this sample, 25% of the data were collected at a pre-tilt stage of 30º to compensate for the preferred orientation of longer chromatin assemblies. After two rounds of 2D-classification to separate hypernucleosomes from cell debris, 1,013,534 particles were used to generate 8 ab-initio classes, followed by heterogeneous refinement at a pixel size of 2.1 Å/px, yielding four different classes of hypernucleosomes (Supp. Fig. 8). After heterogeneous refinement, each class was used for a non-uniform refinement. Following refinement, particles were re-extracted at the original pixel size of 1.053 Å/px. Ab-initio reconstruction into several classes, following a heterogeneous refinement or ab-initio reconstruction with subsequent 3D-classification was performed (Supp. Fig. 8). Each hypernucleosome class was used separately for reference-based motion correction, followed by a final non-uniform refinement, yielding six different hypernucleosome structures, wrapping between 90 - 240 bp of DNA by 3-8 histone dimers, with resolutions up to 3.56 Å (Supp. Fig. 8,10).
***ΔhtkB log-phase dataset*:** 2 rounds of 2D-classifications of two separate template pickers and blob picker particles resulted in 971,195 particles used for ab-initio reconstruction into eight different classes, followed by heterogeneous refinement (Supp Fig. 9). Resulting hypernucleosomes classes were used separately for ab-initio reconstruction at the original pixel size of 1.053 Å/px, followed again by a heterogeneous refinement. After a non-uniform refinement, reference-based motion correction, and a final non-uniform refinement, two hypernucleosome classes were identified with resolutions reaching up to 3.88 Å, containing 3 and 4 dimers, wrapping 90 and 120 bp of DNA (Supp. Fig. 9).

#### Model Fitting and Refinement

First, we have built the models for the best-resolved cryo-EM SPA maps (3.56 Å resolution, log-phase consensus map (5-6 dimer), KU216 SPA dataset; and 3.88 Å resolution, log-phase 4-dimer map, *ΔhtkB* SPA dataset). Starting model for DNA was taken from the closed hypernucleosome structure[17] (PDB 9QV5), and the sequence was substituted for a *T. kodakarensis* gene (Q9Y8I1 UniProt ID). Since the data originate from native chromatin, the underlying DNA sequence is averaged out; to show the difference from the artificial DNA sequences in the literature, we chose a native genomic sequence. Next, HTkA histone dimer models were generated via AlphaFold3[53], and placed into the EM maps. Next, an ISOLDE[54] run in ChimeraX[51] was performed to fit the model into the map and resolve clashes, Ramachandran outliers, and poor rotamers. Next, local adjustments and corrections of the model were made in Coot[55]. Final model quality was assessed via Molprobity[56] and in Coot. For visualization of different side chains of HTkA and HTkB (Fig. 4), residues were mutated in Coot, and a local real-space refinement was performed.

### STA processing workflow
### KU216 stationary phase

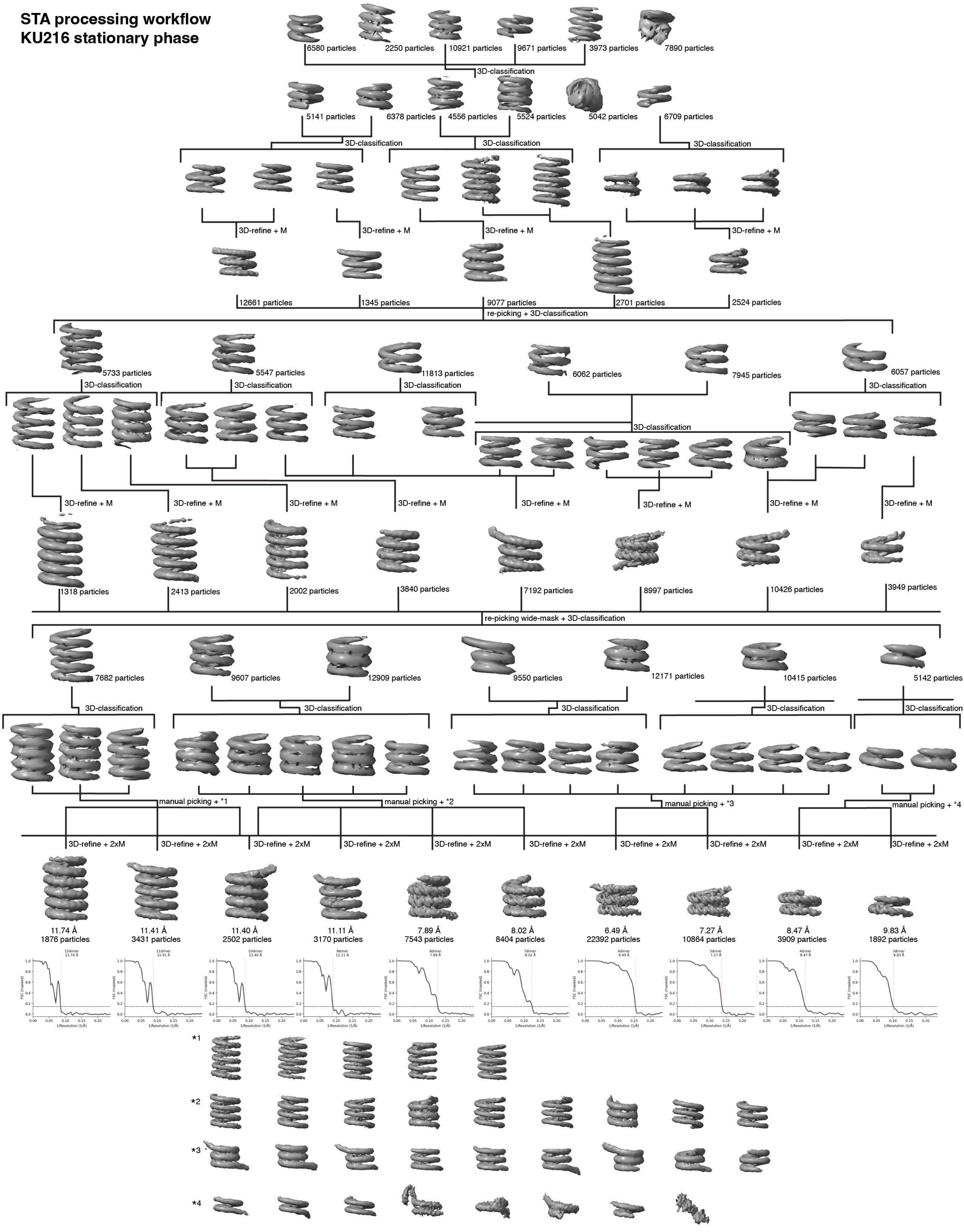

**Supplementary Figure 1:** Cryo-ET subtomogram averaging data processing scheme for KU216 stationary data.

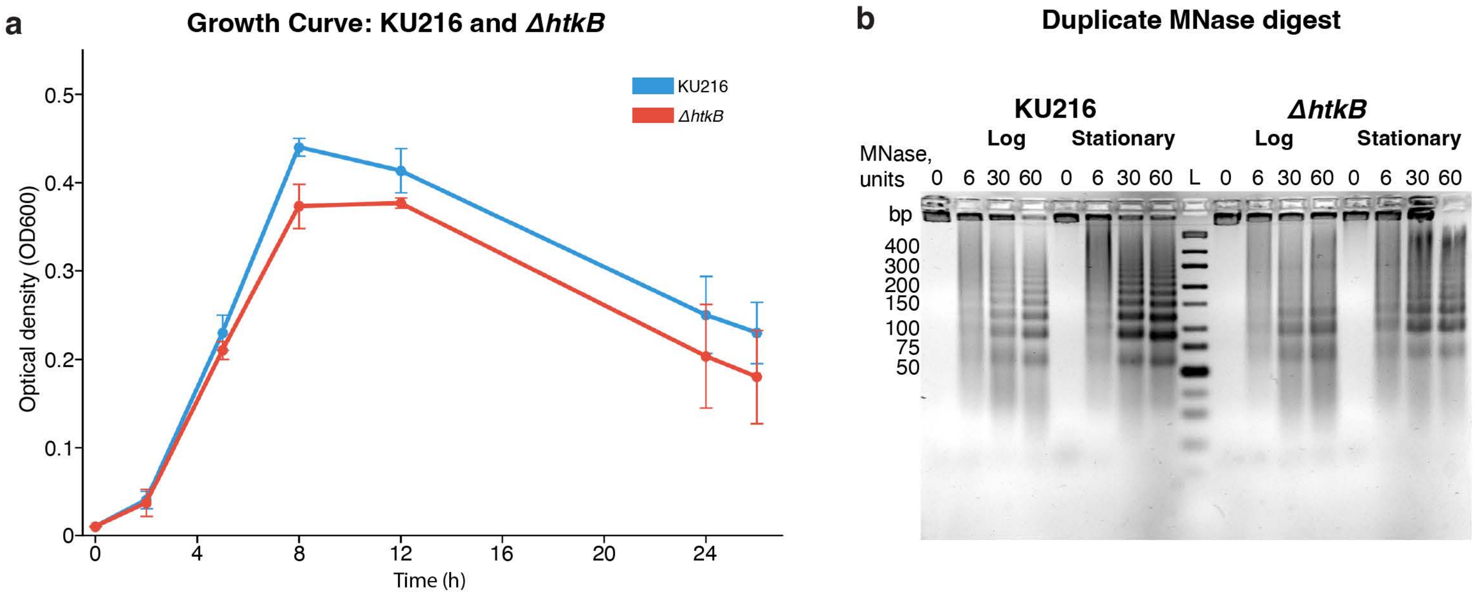


**Supplementary Figure 2:** a. Growth curves for KU216, and ΔhtkB strains of T. kodakarensis. b. Independent biological replicate of the MNase digest experiment (see Fig. 2)

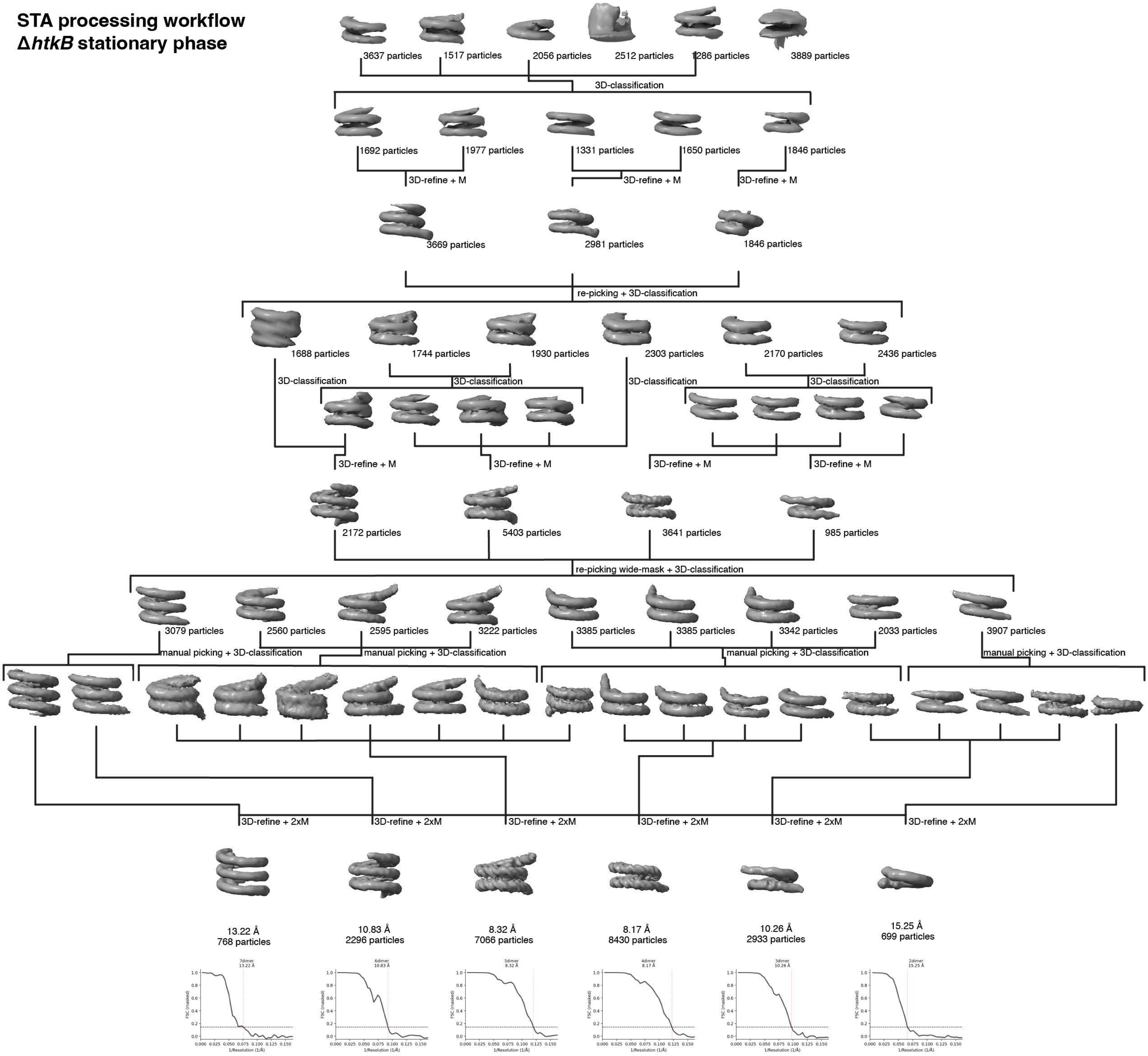


**Supplementary Figure 3:** Subtomogram averaging data processing scheme for Δ*htkB* stationary data. Final subtomogram averages and FSC-curves are displayed at the bottom of the scheme.

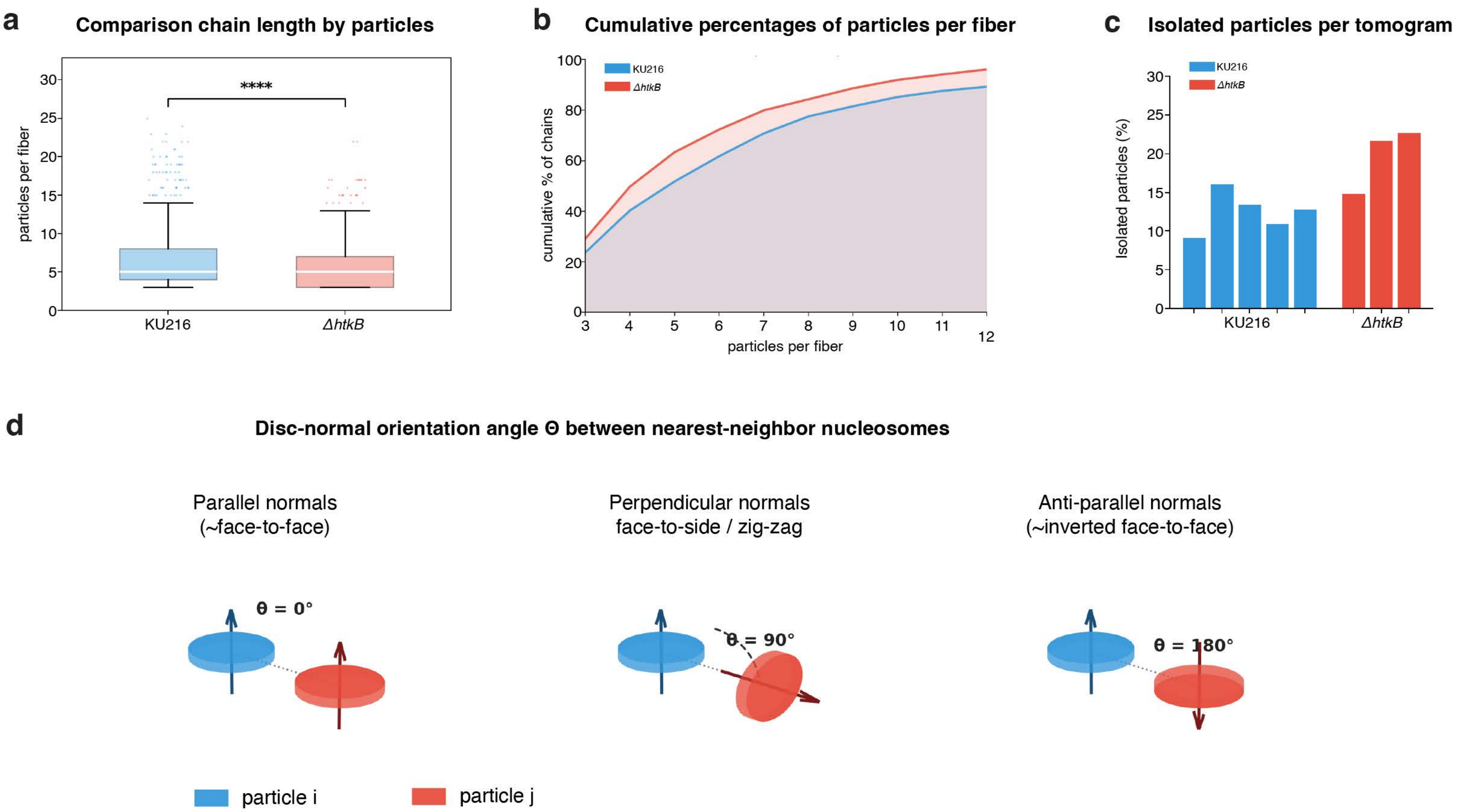


**Supplementary Figure 4**: Chromatin geometry analysis. a. Comparison of chain length by particle counts between KU216 stationary and Δ*htkB* stationary. **** indicates p-value <0.0001 determined by the two-sided Mann-Whitney U test (unpaired). Chains were only considered as chromatin fibers if they consisted of 3 or more particles and the surface-to-surface distance between neighboring particles was 10 nm or less. b. Cumulative distribution of particles per chromatin fiber. c. Percentages of isolated particles in individual tomograms in KU216 (5 tomograms) and Δ*htkB* datasets (3 tomograms). d. Schematic explanation of the relative orientation calculations used for Fig.3b.

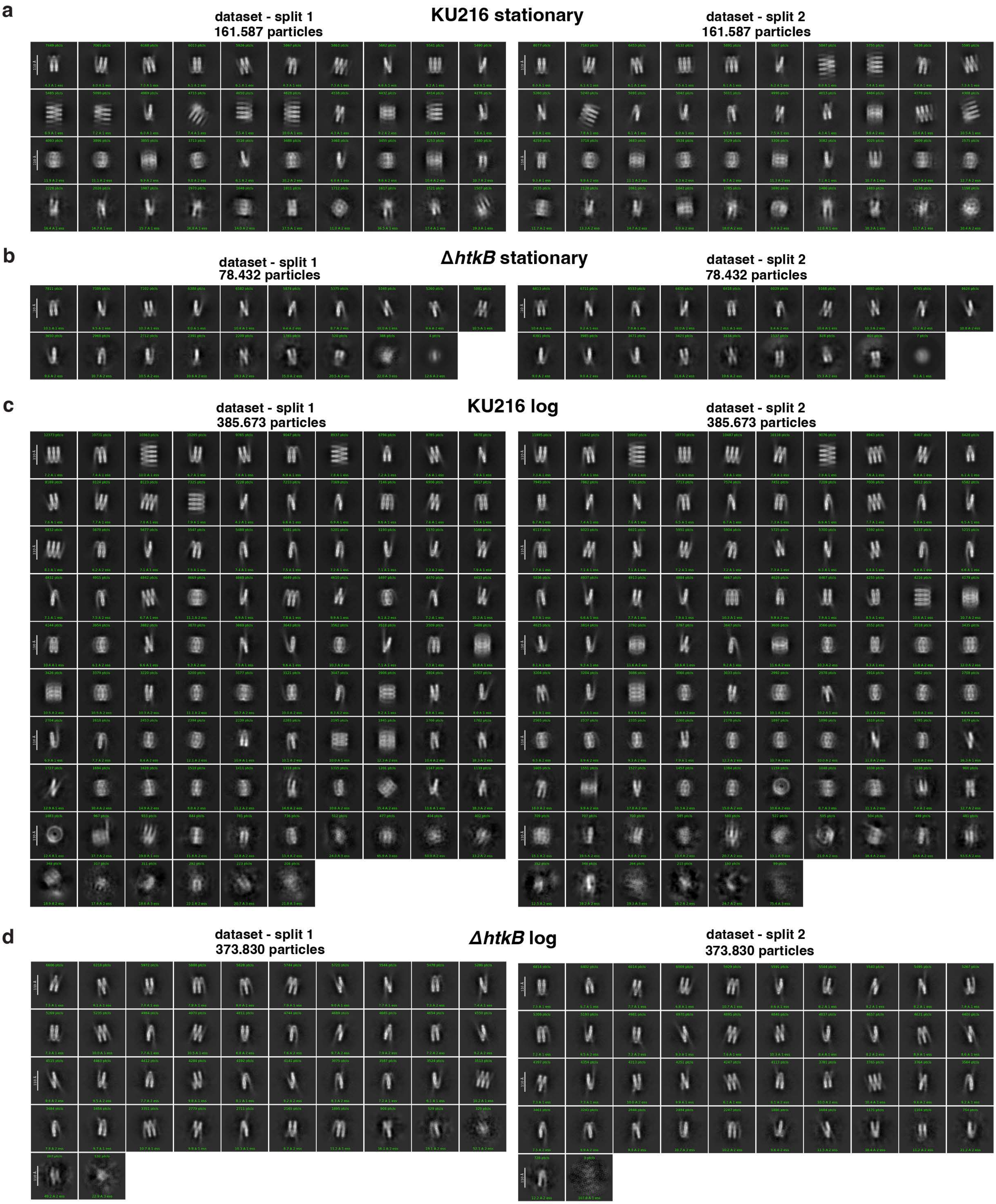


**Supplementary Figure 5:** Selected side views from SPA 2D-classification used for quantification of hypernucleosome size, with each particle set being split into two halves randomly. a. KU216 stationary dataset. b. Δ*htkB* stationary dataset. c. KU216 stationary dataset. d. Δ*htkB* log dataset

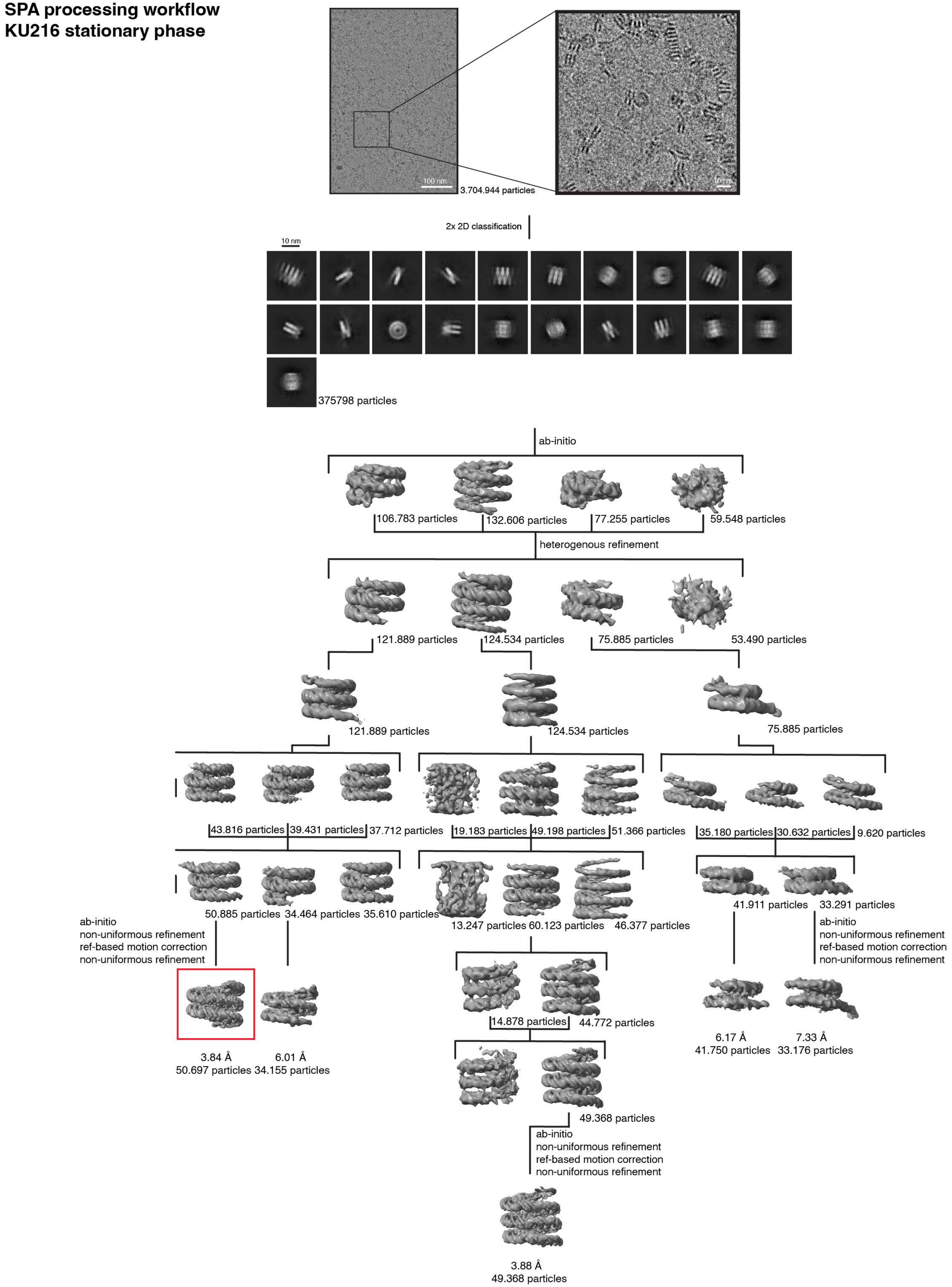


**Supplementary Figure 6:** SPA data processing scheme for KU216 stationary dataset. EM map marked with a red box was used for local resolution estimation in Supp. Fig. 10a.

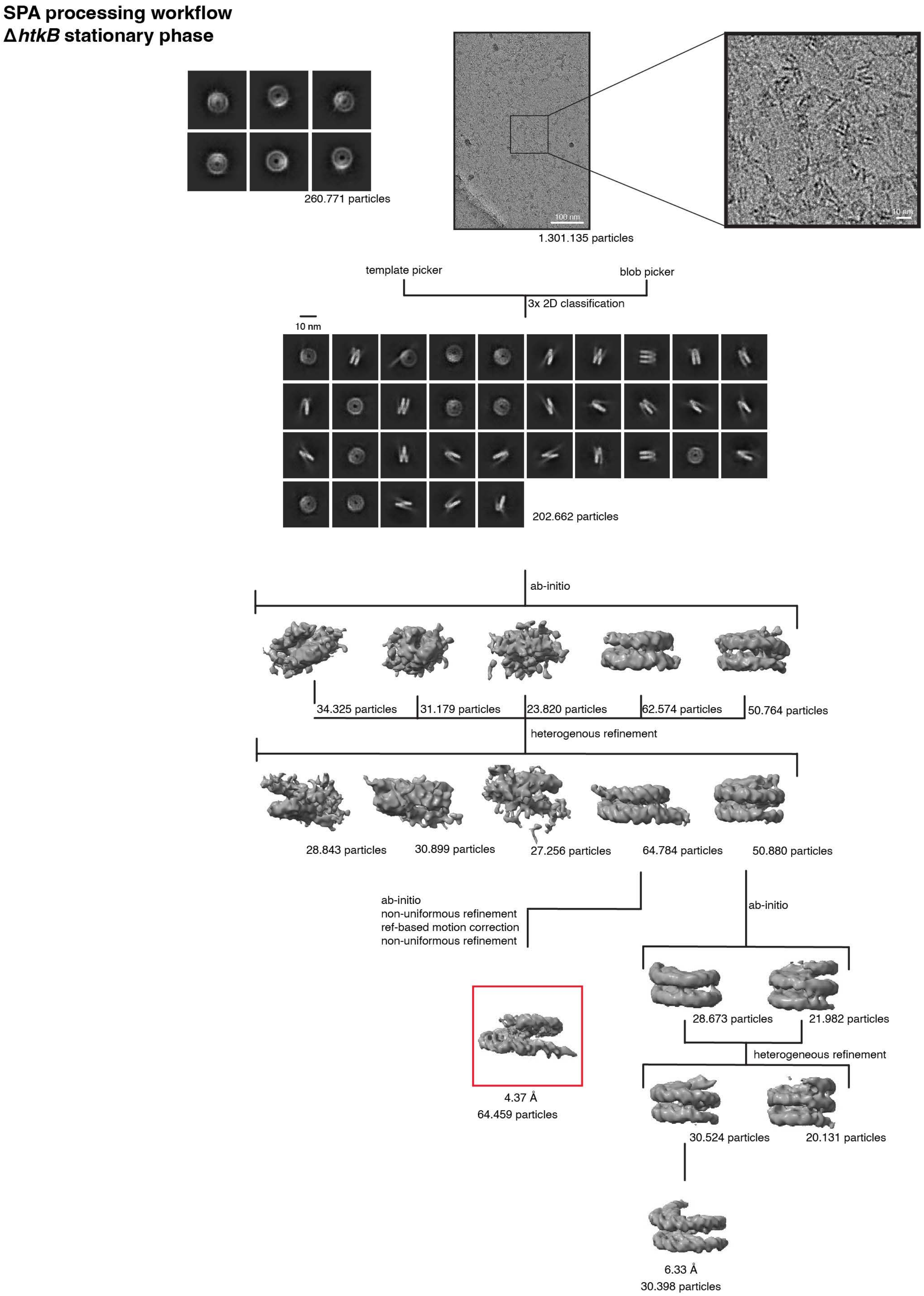


**Supplementary Figure 7:** SPA data processing scheme for Δ*htkB* stationary dataset. EM map marked with a red box was used for local resolution estimation in Supp. Fig. 10.

### SPA processing workflow
### KU216 log phase

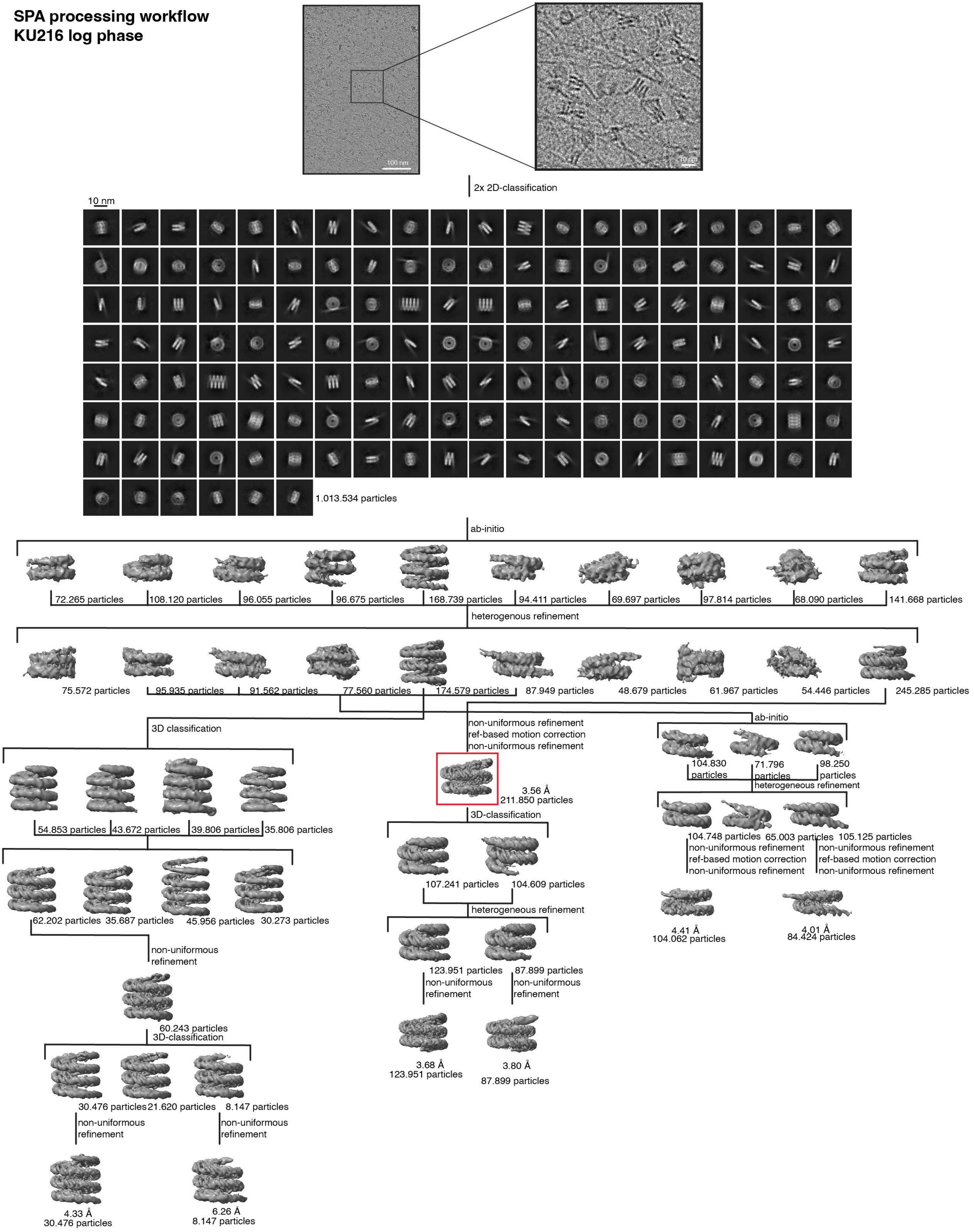

**Supplementary Figure 8:** SPA data processing scheme for KU216 log dataset. EM map marked with a red box was used for local resolution estimation in the Supp. Fig. 10.

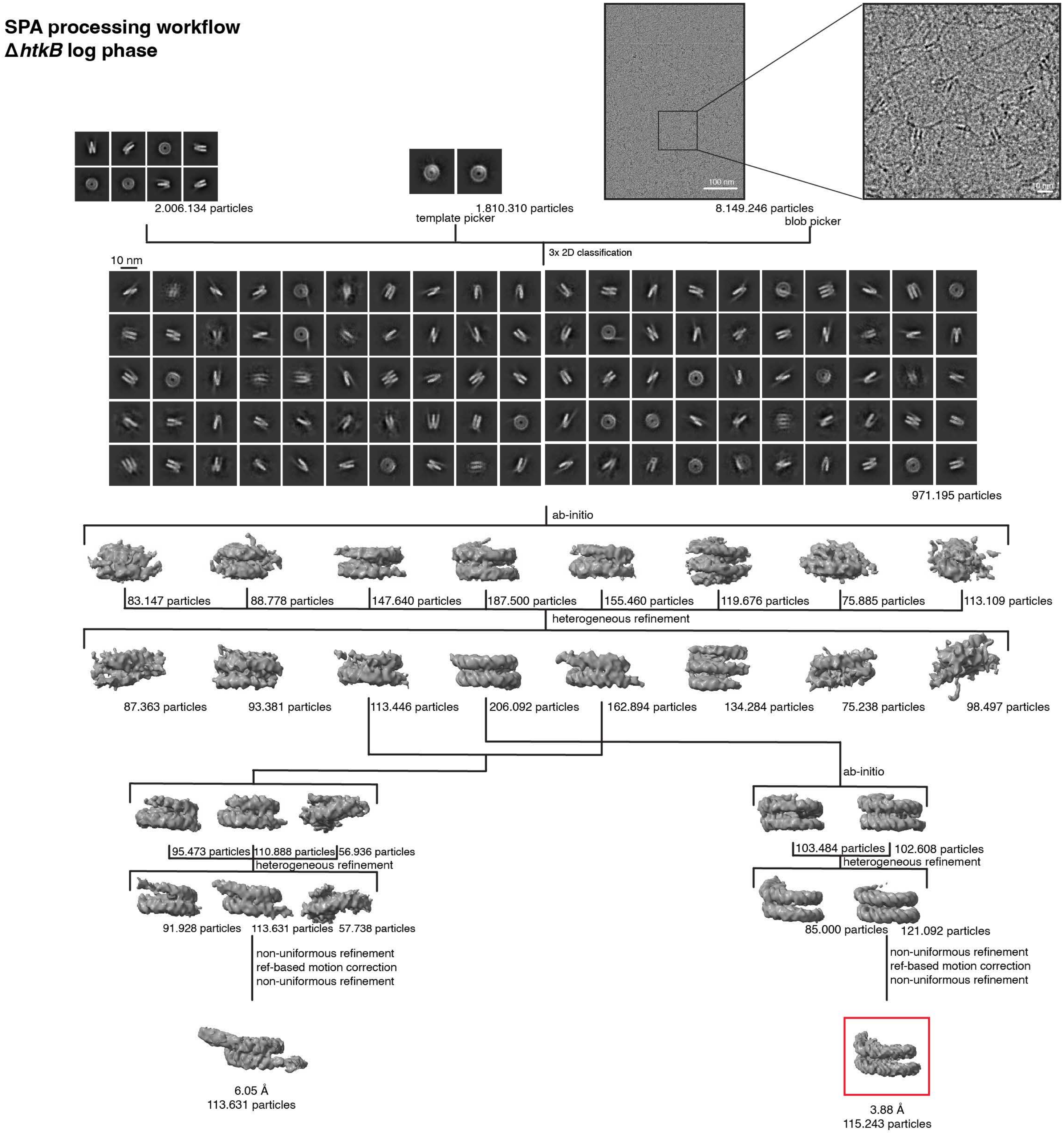


**Supplementary Figure 9:** SPA data processing scheme for Δ*htkB* log dataset. EM map marked with a red box was used for local resolution estimation in Supp. Figure 10.

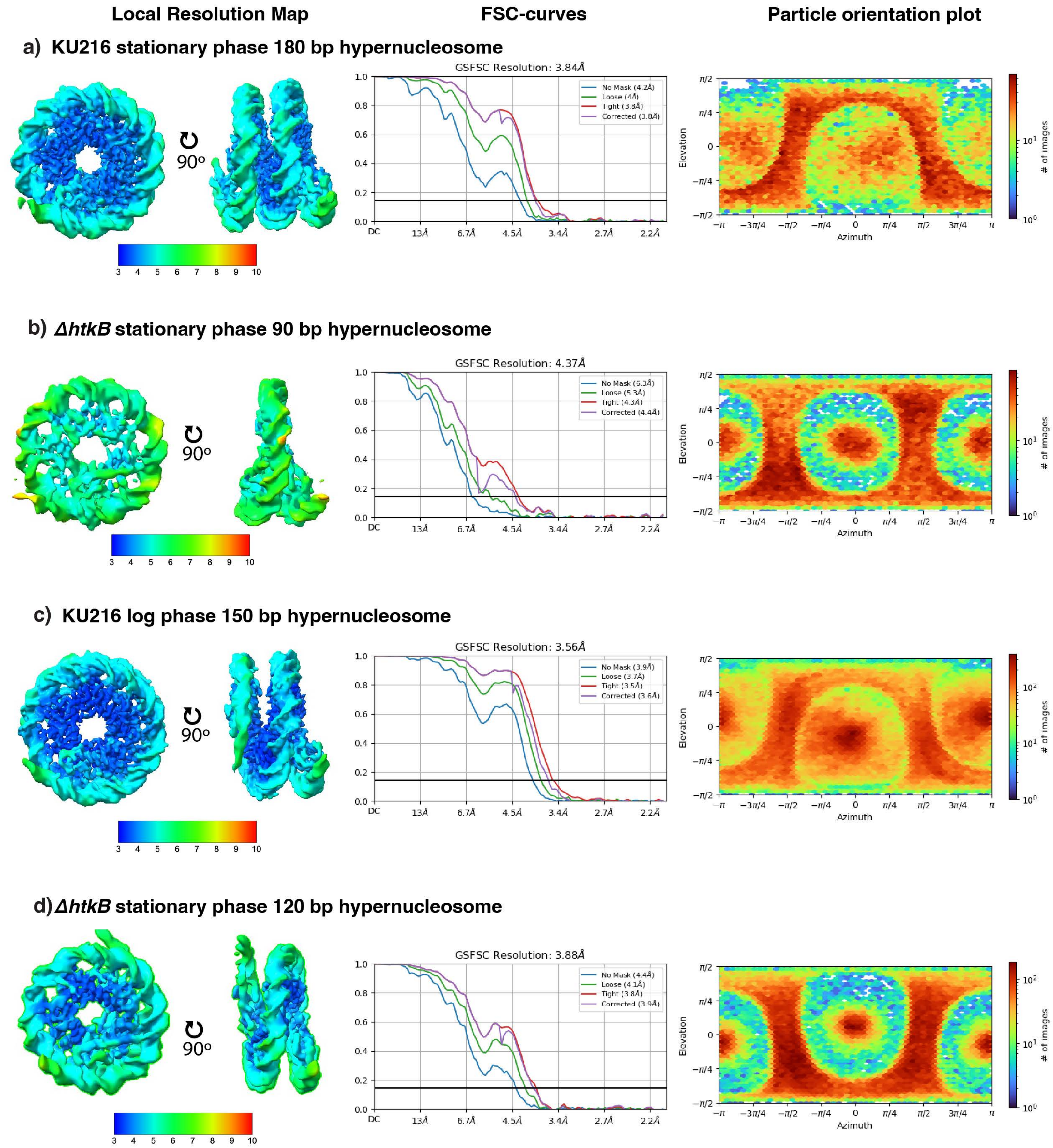


**Supplementary Figure 10:** For the highest resolved map of each of the four SPA datasets: Local resolution estimation, FSC-curves, particle orientation discributions. a. KU216 stationary phase 180 bp hypernucleosome. b. Δ*htkB* stationary phase 90 bp hypernucleosome. c. KU216 log phase 150 bp hypernucleosome. d. Δ*htkB* stationary phase 120 bp hypernucleosome
